# Two distinct mechanisms of Plexin A function in *Drosophila* optic lobe lamination and morphogenesis

**DOI:** 10.1101/2023.08.07.552282

**Authors:** Maria E. Bustillo, Jessica Douthit, Sergio Astigarraga, Jessica E. Treisman

## Abstract

Visual circuit development is characterized by subdivision of neuropils into layers that house distinct sets of synaptic connections. We find that in the *Drosophila* medulla, this layered organization depends on the axon guidance regulator Plexin A. In *plexin A* null mutants, synaptic layers of the medulla neuropil and arborizations of individual neurons are wider and less distinct than in controls. Analysis of Semaphorin function indicates that Semaphorin 1a, provided by cells that include Tm5 neurons, is the primary partner for Plexin A in medulla lamination. Removal of the cytoplasmic domain of endogenous Plexin A does not disrupt the formation of medulla layers; however, both null and cytoplasmic domain deletion mutations of *plexin A* result in an altered overall shape of the medulla neuropil. These data suggest that Plexin A acts as a receptor to mediate morphogenesis of the medulla neuropil, and as a ligand for Semaphorin 1a to subdivide it into layers. Its two independent functions illustrate how a few guidance molecules can organize complex brain structures by each playing multiple roles.

**Summary statement:** The axon guidance molecule Plexin A has two functions in *Drosophila* medulla development; morphogenesis of the neuropil requires its cytoplasmic domain, but establishing synaptic layers through Semaphorin 1a does not.

## Introduction

During the development of the nervous system, cell-cell interactions driven by cell-surface molecules determine the formation of synaptic connections. Such connections are often concentrated in synaptically dense regions called neuropils. Spatial segregation of the synaptic connections between different neuronal types, for instance in layers or nuclei, simplifies the process of growth cone navigation to the correct target cells (Sanes and Zipursky, 2010). Synaptic laminae are found in many regions of both vertebrate and invertebrate nervous systems, including the *Drosophila* optic lobe (Millard and Pecot, 2018; Sanes and Zipursky, 2010), the vertebrate retina (Baier, 2013) and the mammalian cortex and cerebellum (Guy and Staiger, 2017). These layers reflect the segregation of neurites of distinct cell types and the formation of localized connections with specific synaptic partners.

Several mechanisms that establish a laminated organization of brain structures have been identified in the vertebrate retina. Synaptic laminae in the inner plexiform layer of the chick retina depend on differential localization of members of the immunoglobulin (Ig) domain-containing family of proteins – two Sidekicks, five Contactins, and two Dscams - in non-overlapping groups of bipolar, amacrine and ganglion cells (Yamagata and Sanes, 2008; Yamagata and Sanes, 2012). Homophilic interactions between these molecules are necessary and sufficient to direct neurites that express the same Ig protein to the same layer (Yamagata et al., 2002). Another mechanism used to organize sublaminae in the mouse inner plexiform layer is the complementary localization of receptor-ligand pairs. For example, localization of Semaphorin 6A in ON laminae, Plexin A4 in OFF laminae, and Plexin A2 in both directs the targeting of multiple neuronal types and establishes the stratification of the motion detection pathway via repulsive signaling (Matsuoka et al., 2011; Sun et al., 2013). Additional insights into layer formation may be revealed by studying a simpler model system with fewer gene duplications.

The *Drosophila* medulla, the largest of the four neuropils in the optic lobe, is a laminated structure that resembles the inner plexiform layer of the vertebrate retina (Sanes and Zipursky, 2010). It represents many highly stereotyped and segregated neurite projections from numerous cell types, and its ten layers emerge over the course of pupal development (Ngo et al., 2017). As in the chick retina, *Drosophila* Sidekick (Sdk) is present in distinct layers of the medulla neuropil; however, loss of *sdk* does not disrupt medulla layers (Astigarraga et al., 2018). Lamination in the fly medulla also requires complementary localization of Plexins and Semaphorins. Lamina neurons are interneurons that receive input from motion-sensitive photoreceptors and form output synapses in specific layers of the outer medulla neuropil. Repulsive interactions between Semaphorin 1a (Sema1a) expressed in all lamina neurons and Plexin A (PlexA) expressed by tangential neurons in the M7 layer of the medulla prevents these lamina neurons from projecting into the inner medulla (Pecot et al., 2013). Thus, the subdivision of neuropils into functional synaptic laminae is a highly conserved organizational strategy that can be accomplished by both adhesive and repulsive mechanisms.

PlexA is one of two *Drosophila* members of the conserved Plexin family of transmembrane signaling proteins. Plexins interact with their canonical partners, Semaphorins, through the extracellular Sema domains of both proteins (Andermatt et al., 2014; Janssen et al., 2010; Liu et al., 2010; Nogi et al., 2010; Winberg et al., 1998). Plexin A proteins are well established as receptors for class I transmembrane Semaphorins during axon guidance in vertebrates and invertebrates (Hung et al., 2010; Negishi et al., 2005; Tamagnone et al., 1999), and signal through an intracellular RasGAP domain as well as interacting with other downstream effectors (Ayoob et al., 2004; Hung et al., 2010; Pascoe et al., 2015). Overexpression studies suggest that Plexin A proteins can also act as ligands that activate a Semaphorin 1 reverse signaling pathway to remodel the actin cytoskeleton (Andermatt et al., 2014; Jeong et al., 2012; Jeong et al., 2017; Sun et al., 2015; Toyofuku et al., 2004; Yu et al., 2010).

Here, we show that PlexA is required during pupal development to establish synaptic layering in the *Drosophila* medulla. Loss of *plexA* disrupts neuronal projections throughout the medulla, resulting in global disorganization of medulla layers. We use CRISPR modification of the endogenous *plexA* gene to show that PlexA mediates layering independently of its cytoplasmic domain, likely serving as a ligand for Sema1a in specific medulla neuron types such as the Tm5 neurons. In contrast, PlexA requires its cytoplasmic domain to establish the overall shape of the medulla neuropil during pupal development. We propose that PlexA has dual functions during medulla development, as a receptor to control neuropil morphogenesis, and as a ligand for Sema1a to establish neurite segregation that gives rise to a laminated medulla architecture.

## Results

### Loss of *plexA* disrupts R7 axon targeting

Targeting of color-sensitive R7 and R8 photoreceptor axons to distinct layers of the *Drosophila* medulla is a powerful model with which to investigate the organization of synaptic layers. Several studies have identified factors that act within the photoreceptors themselves to enable R8 to terminate in the M3 layer of the medulla and R7 to project beyond it to the M6 layer (Astigarraga et al., 2010; Plazaola-Sasieta et al., 2017). However, fewer of the molecular cues that photoreceptor axons recognize within the medulla to allow them to target these specific layers are known (Timofeev et al., 2012). To identify such cues, we carried out an RNAi screen in which we knocked down genes predicted to encode cell-surface and secreted molecules (Kurusu et al., 2008) with a combination of three GAL4 drivers (*apterous* (*ap*)*-GAL4, Distal-less* (*Dll*)*-GAL4, and eyeless* (*ey^OK107^*)*-GAL4*) that together are expressed in a large proportion of medulla neurons (Li et al., 2013; Morante and Desplan, 2008)(Fig. S1A). We found that knocking down *plexA* caused R7 axons to terminate prematurely in the same layer as R8 axons in the anterior medulla (Fig. 1A, B). A similar phenotype was observed using a second RNAi line targeting *plexA* (Fig. 1C, Fig. S1C), indicating that it was not due to an off-target effect.

**Figure 1.**
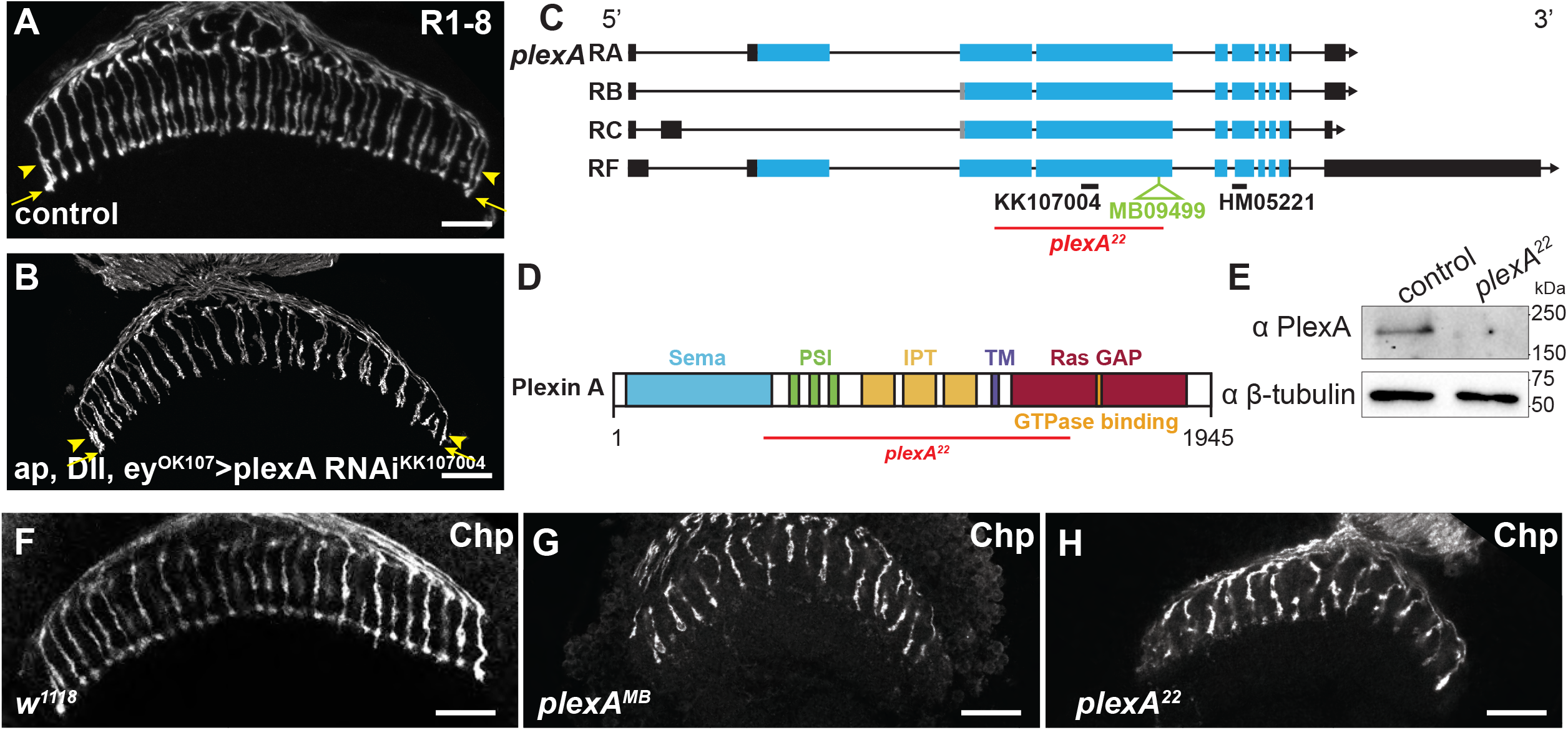
*plexA* is required for R7 axon targeting. **(A, B)** Adult head cryosections showing *w^1118^* control **(A)** and knockdown of *plexA* in medulla neurons with *ap-GAL4*, *Dll-GAL4* and *ey^OK107^-GAL4* **(B)**. Photoreceptors marked with *gl-lacZ*; M3 layer marked with arrowhead, M6 layer marked with arrow. (**C)** Schematic of PlexA isoforms, denoting site of the Minos insertion in *plexA^MB09499^*(green), the deletion in *plexA^22^* (red), and two RNAi lines (black). **(D)** Schematic of PlexA protein showing the region deleted in *plexA^22^*. **(E)** Western blot confirming the absence of PlexA protein in the *plexA^22^*mutant allele. **(F-H)** Late pupal whole mount *w^1118^* control **(F),** *plexA^MB09499^* **(G)** and *plexA^22^***(H)** mutant medullas showing disruption of photoreceptor axons in *plexA* alleles **(G, H)**. Photoreceptors marked with anti-Chp. Scale bar = 20 µm. Images are taken in a horizontal plane with anterior to the left and posterior to the right in this and all subsequent figures, except when indicated.

PlexA is a transmembrane protein that contains an N-terminal Sema domain, three Plexin, Semaphorin and Integrin (PSI) domains, and three Immunoglobulin-like, Plexin and Transcription factor (IPT) domains in the extracellular region, and an intracellular bipartite RasGAP domain (Fig. 1D). To confirm that *plexA* is required for normal photoreceptor axon projections, we used CRISPR mutagenesis to delete a large region including sequences that encode the transmembrane domain and much of the extracellular domain (Fig. 1C, D), generating a protein null allele, *plexA^22^* (Fig. 1E). We also examined *plexA^MB09499^*(Bellen et al., 2011), a *Minos* transposable element insertion into a coding exon (Fig. 1C). Homozygotes for either mutation die prior to eclosion. We found that in *plexA* homozygous mutant pupae, R7 and R8 projections were disordered and did not target to distinct layers in the medulla (Fig. 1F-H). Additionally, *plexA* mutant photoreceptor axons were misshapen and formed less distinct terminals than controls (Fig. 1G, H).

*plexA* is expressed in the eye as well as the brain, and one of the GAL4 lines we used for our screen, *ey^OK107^-GAL4*, also drives expression in the eye. To test whether these defects were caused by loss of *plexA* in medulla neurons or in photoreceptors, we generated homozygous *plexA* mutant clones in the eye using a duplication of the *plexA* genomic region inserted on the third chromosome. R7 photoreceptor axon projections terminated correctly in the M6 layer, even in very large *plexA* mutant clones generated in a *Minute* background (Fig. S1D), indicating that *plexA* is not required autonomously in photoreceptor cells. Taken together, these data demonstrate that normal photoreceptor axon targeting requires *plexA* function in medulla neurons but not in photoreceptors.

### PlexA is required for normal patterning of medulla neuron projections

During mid-pupal development, R7 cells form synapses with Dm8 interneurons and other post-synaptic partners in the M6 layer (Gao et al., 2008; Kind et al., 2021; Menon et al., 2019). It is possible that mistargeting of photoreceptor axons would preclude these cells from forming synapses, if Dm8 neurites were not able to contact photoreceptor axon terminals. We labelled Dm8 cells with *UAS-myrTomato* driven by *ort^C2b^-GAL4* in *plexA* mutant flies to determine whether R7 axons terminate before reaching these target cells. In control pupae, Dm8 dendrites arborize primarily in the M6 layer, where they make contact with R7 axons (Ting et al., 2014)(Fig. 2A). In *plexA* mutant pupae, Dm8 projections were disordered and extended into a wider region of the medulla, enabling them to contact the mis-projecting R7 axon terminals (Fig. 2B, I). To ascertain whether these contacts contained synaptic components, we expressed a tagged short version of the presynaptic protein Bruchpilot (Brp) in a subset of R7 cells (Fig. S1E, F). Brp puncta were still present in R7 terminals when *plexA* was knocked down in all neurons with *elav-GAL4*, suggesting that synaptogenesis still occurred despite the abnormal position of R7-Dm8 contacts (Fig. S1F). Pan-neuronal knockdown of *plexA* using *elav-*GAL4 disrupted R7 targeting (Fig. S1F) but did not prevent eclosion; the late pupal lethality observed in whole animal null mutants is thus likely due to a non-neuronal developmental role for PlexA.

**Figure 2.**
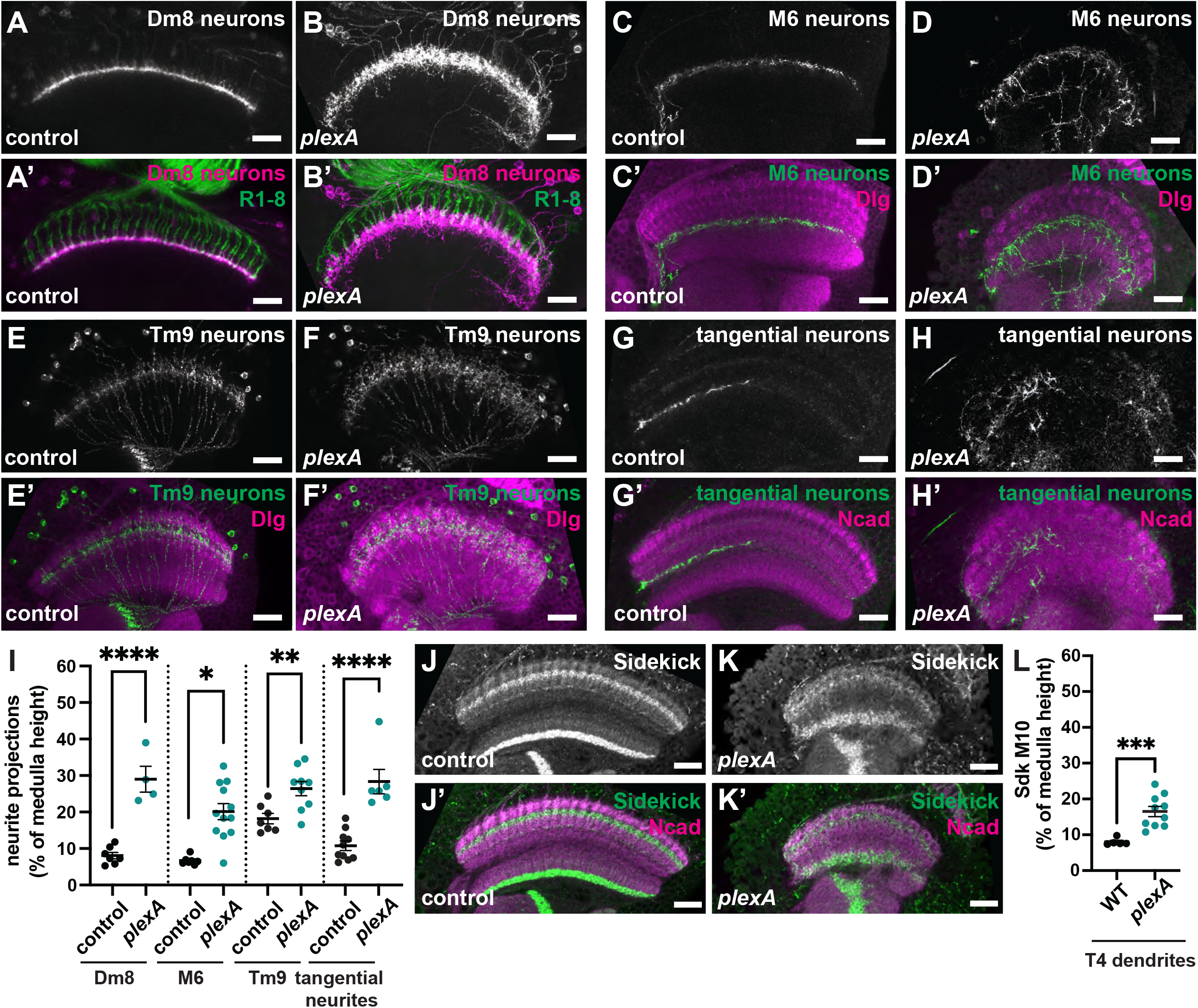
*plexA* is required for normal patterning of medulla neuron projections. **(A-H, J, K)** show 72 h APF brains. **(A, C, E, G, J)** *w^1118^* control; **(B, D, F, H, K)** *plexA^22^/plexA^MB09499^*. **(A, B)** Visualization of Dm8 neurons with *ort^C2b^-*GAL4 driving *UAS-myrTomato*. **(A’, B’)** Tomato shown in magenta and Chp in green. **(C-F)** Visualization of M6 local neurons with *30A*-GAL4 **(C, D)** and Tm9 neurons with GMR*24C08*-GAL4 **(E, F)** driving *UAS-myrTomato*. **(C’-F’)** Tomato shown in green and Dlg in magenta. **(G, H)** Tangential neurons visualized with *GMR35A02*-GAL4 driving *UAS-myrTomato*. **(G’, H’)** Tomato shown in green and Ncad in magenta. Scale bar = 20 µm. **(I)** Quantification of proximal-distal extent of neurites in controls and *plexA^22^/plexA^MB09499^* mutants, plotted as a percentage of medulla height. N = 7 control, 4 *plexA* Dm8 samples; 7 control, 12 *plexA* M6 samples; 7 control, 9 *plexA* Tm9 samples; 10 control, 6 *plexA* tangential neuron samples. **(J, K)** Brains stained for Sdk to label T4 dendrites in the M10 layer. **(J’, K’)** Sdk shown in green and Ncad in magenta. **(L)** Quantification of proximal-distal extent of Sdk staining in the M10 layer of *w^1118^* control and *plexA^22^* brains as a percentage of medulla height. N = 5 WT, 10 *plexA^22^*. Measurements were compared via unpaired t-test. Error bars show mean ± SEM. *, P ≤ 0.05; **, P ≤ 0.01; ***, P ≤ 0.001; ****, P ≤ 0.0001.

The effect of *plexA* on Dm8 dendrite projections suggested that rather than specifically guiding R7 photoreceptors, PlexA might play a more general role in establishing medulla neuron projection morphology. To examine the effect of loss of *plexA* on other medulla neurons, we labelled local interneurons in the M6 layer, Tm9 projection neurons, and tangential neurons in the M7 layer with *UAS-myrTomato* driven by *30A-GAL4* (Chin et al., 2014)*, GMR24C08-GAL4* (Fisher et al., 2015), and *GMR35A02-GAL4* (Jenett et al., 2012) respectively. The neurites of *30A-GAL4*-expressing interneurons appeared tightly apposed in the M6 layer in wildtype brains (Fig. 2C). In *plexA* mutants, these interneurons instead extended projections through many layers of the medulla (Fig. 2D, I). Tm9 neurons have dendritic arborizations in layer M3, and their axons project through the medulla in highly regular parallel columns, terminating in lobula layer 1 (Fischbach and Dittrich, 1989) (Fig. 2E). Tm9 projections were disordered in *plexA* mutants, with the M3 arborizations expanding to cover a larger portion of the medulla (Fig. 2F, I). Medulla tangential neurons send processes horizontally across the medulla in the M7 layer (Fig. 2G). We found that in *plexA* mutant flies, tangential neuron projections were not confined to this layer, but wandered extensively throughout the proximal medulla (Fig. 2H, I). Finally, we used an anti-Sdk antibody to label the dendrites of T4 neurons in layer M10 (Astigarraga et al., 2018) and observed that Sdk staining in this layer occupied a larger proportion of the width of the medulla in *plexA* mutants than in controls (Fig. 2J-L). Loss of *plexA* thus disrupts the projection patterns of neurites that project in several layers and belong to multiple medulla neuron cell types.

### PlexA is required to establish a layered architecture in the medulla

To characterize how loss of *plexA* affects the overall organization of the medulla, we stained *plexA* mutant pupal brains 72 h after puparium formation (APF) for the postsynaptic protein Discs-large (Dlg). Visualization of Dlg in wild-type optic lobes showed a consistent pattern that varied in intensity between individual medulla layers, with notably high intensity in layers M2, the proximal half of M3, M5, M9, and M10 (Chin et al., 2014)(Fig. 3A). Dlg staining in *plexA mutants* was not as tightly restricted to distinct layers as in wild-type pupae (Fig. 3B), and the peaks and valleys of Dlg intensity that characterize the wild-type medulla were flattened (Fig. 3C). The neuronal cadherin family member, N-cadherin (Ncad) was present throughout much of the wild-type medulla neuropil, with three distinct regions of low Ncad intensity in layers M3, M5 and the proximal half of M7 (Fig. 3D, F). In *plexA* mutants, Ncad staining was more uniform across the medulla (Fig. 3E, F). We quantified this difference by measuring the sinuosity of the intensity curves (their difference in length from a straight line connecting the endpoints). The sinuosity of both measurements was significantly reduced in *plexA* mutants (Fig. 3G). Most of the differences in intensity in *plexA* mutants affected the medial medulla, with layers M1, M2, and M10 appearing relatively normal (Fig. 3C, F). The poorly defined layers in the *plexA* mutant medulla may explain the mistargeting of R7 photoreceptor axons.

**Figure 3.**
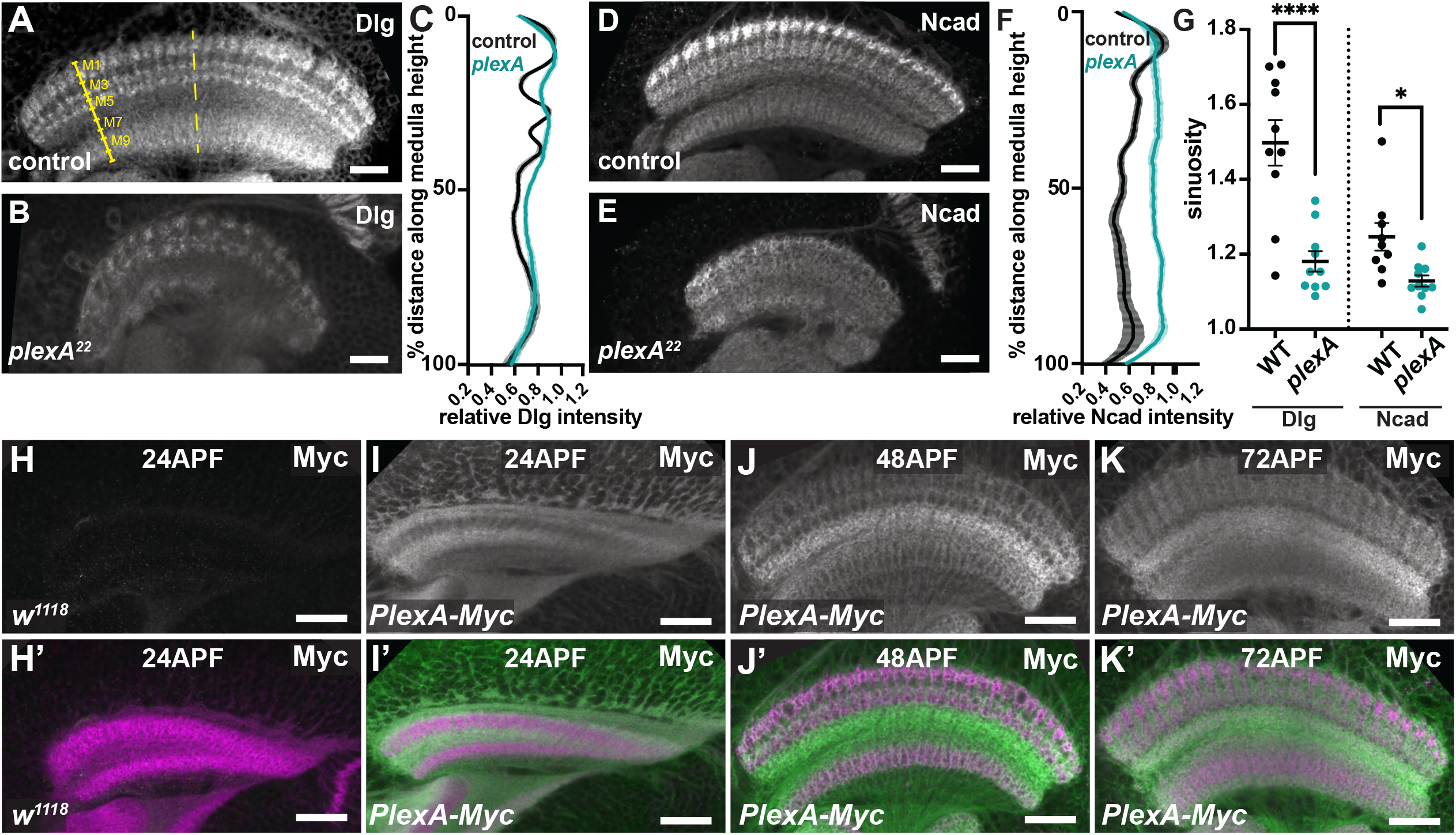
p*lexA* is required for global medulla lamination. **(A, B, D, E)** Control *w^1118^***(A, D)** and *plexA^22^* mutant **(B, E)** 72 h APF brains stained for Dlg **(A, B)**, and Ncad **(D, E)**. Synaptic layers M1-10 are labeled based on Dlg intensity shown in yellow in **(A)**, as in Chin et al., 2014. **(C, F)** Line intensity traces of Dlg **(C)** and Ncad **(F)** staining in control (black) and *plexA^22^* mutant medullas (teal) over the height of the medulla, as shown in yellow dashed line in **(A)**. Distance is normalized as a percentage of medulla height; staining intensity at each position is normalized to the highest intensity value for that condition. Solid line represents the average of multiple samples, error bars represent SEM. N = 10 brains/ genotype. **(G)** shows measurements of the sinuosity of these Dlg and Ncad curves (ratio of the curve length to that of a straight line connecting the endpoints). N = 10 WT and *plexA^22^* Dlg samples; 9 WT and 10 *plexA^22^* Ncad samples. Measurements were compared via unpaired t-test. Error bars show mean ± SEM. *, P ≤ 0.05; ****, P ≤ 0.0001. **(H-K)** PlexA-Myc staining with anti-Myc showing PlexA protein in the medulla and its enrichment in the M7 layer at 24 h APF **(I)**, 48 h APF **(J)**, and 72 h APF **(K)**. **(H)** shows that Myc staining was undetectable in a *w^1118^* 24 h APF brain. **(H’-K’)** Myc shown in green, neuropil labeled with Ncad in magenta. Scale bars = 20 µm.

We next investigated where PlexA acts to promote medulla layering and photoreceptor axon targeting. Expressing *plexA* RNAi in all neurons with *elav-GAL4* resulted in shortened and disorganized photoreceptor axon projections, similar to *plexA* mutants (Fig. S1E, F), indicating that *plexA* acts in neurons. In the pupal medulla, PlexA protein is enriched in the developing serpentine layer, M7 (Fig. 3H-K, Fig. S2A-D), where it colocalizes with the processes of medulla tangential neurons (Pecot et al., 2013). Medulla neurons largely arise from two neuronal progenitor populations: the main outer proliferation center (OPC), and the tips of the OPC (tOPC) (Bertet et al., 2014; Li et al., 2013). Neuroblasts that form the main OPC express a temporal series of transcription factors beginning with Homothorax (Hth) and go on to generate most of the neurons in the medulla (Li et al., 2013). However, neuroblasts in the tOPC, which express a distinct series of transcription factors beginning with Distal-less (Dll), give rise to medulla tangential neurons among other cell types (Bertet et al., 2014). To determine which neuroblast lineage is responsible for PlexA expression, we expressed *UAS-Cas9* and *plexA* sgRNAs with *hth-GAL4* and *Dll-GAL4* drivers in order to mutate *plexA* in each population of neuroblasts by somatic CRISPR. Loss of *plexA* in neurons of the *hth-GAL4* lineage did not significantly affect photoreceptor axon targeting or Ncad staining (Fig. S2F, H). In contrast, loss of *plexA* in neurons of the *Dll-GAL4* lineage resulted in shortened and disorganized R7 and R8 axon projections as well as less distinct Ncad layering when compared to control brains (Fig. S2E-H), suggesting that *plexA* is required in neurons derived from tOPC neuroblasts. The weaker phenotype compared to *plexA* mutants could reflect mosaic rather than complete deletion of the *plexA* gene, and/or a contribution of additional neurons to PlexA function. Although PlexA was enriched in the M7 layer during mid- and late pupal development, some PlexA protein was detected throughout the medulla neuropil (Fig. S2B, C), and could be clearly visualized using a Myc-tagged form of endogenous PlexA (Fig. 3H-K). These data indicate that PlexA expressed by M7 tangential neurons formed from tOPC neuroblasts contributes to the establishment of layers in the medulla neuropil, but these cells may not be the only source of PlexA.

### PlexA establishes medulla layering through Sema1a

Interactions between PlexA and the Semaphorin family of proteins are important for vertebrate and invertebrate axon guidance (Negishi, Oinuma and Katoh, 2005; Tamagnone et al., 1999) as well as other developmental processes such as epithelial remodeling following wound healing (Yoo et al., 2016) and collective cell migration (Stedden et al., 2019). Plexins and Semaphorins interact through their Sema domains, which are highly conserved between vertebrates and invertebrates (Janssen et al., 2010; Winberg et al., 1998). Interestingly, the bulk of the Sema domain of PlexA is encoded by an exon that is missing from splice isoforms RB and RC (Fig. 1C). To determine whether the Sema domain is necessary for the function of PlexA in medulla layer formation, we specifically mutated this exon in the *plexA* gene by CRISPR (*plexA*Δ*Sema*) and stained brains for Ncad and Sdk to visualize neuropil layers. We found that in *plexA*Δ*Sema* mutants, photoreceptor targeting was disrupted and the medulla layers labeled by Ncad and Dlg were wider and more diffuse (Fig. S3A, B), similar to *plexA* null mutants. This suggests that isoforms containing the Sema domain (RA and RF) are necessary to pattern the medulla layers. Western blotting indicated the presence of low levels of PlexAΔSema protein in larval brains (Fig. S3C).

Three *Drosophila* Semaphorins - Sema1a, Sema1b, and Sema5c - interact with PlexA (Stedden et al., 2019; Winberg et al., 1998) and are expressed in the pupal optic lobe throughout development (Kurmangaliyev et al., 2020; Ozel et al., 2021). Sema1a protein is present within the medulla as early as 12 h APF, but is excluded from the putative M7 layer (Pecot et al., 2013). To examine whether PlexA establishes medulla layering via interactions with Semaphorin family members, we stained *sema1a^P1^* (Yu et al., 1998), *sema1b^KO^* (Wittes and Schüpbach, 2019), and *sema5c^K175^* (Stedden et al., 2019) mutant late pupal medullas for Dlg and Ncad (Fig. 4A, B, Fig. S3D, E). Neither *sema1b* nor *sema5c* disrupted the establishment of medulla layers visualized by Ncad and Dlg staining (Fig. S3D, E). Loss of *sema1a* decreased the sinuosity of Ncad and Dlg measurements, indicating that layers are less distinct in *sema1a* mutants (Fig. 4C-E). Examination of the intensity profiles of Ncad and Dlg immunofluorescence showed that loss of *sema1a* reduced the definition of Ncad layers primarily in the medial medulla (between the 20th – 70th percentiles of medulla height) (Fig. 4B’, D), and distal Dlg layers appeared better defined than in *plexA* mutants (compare Fig. 4B, C with 3B, C). We also found that in *sema1a* mutant pupae, tangential neurons extended neurite projections outside layer M7 of the medulla rather than forming a tight bundle as in control brains (Fig. 4F-H).

**Figure 4.**
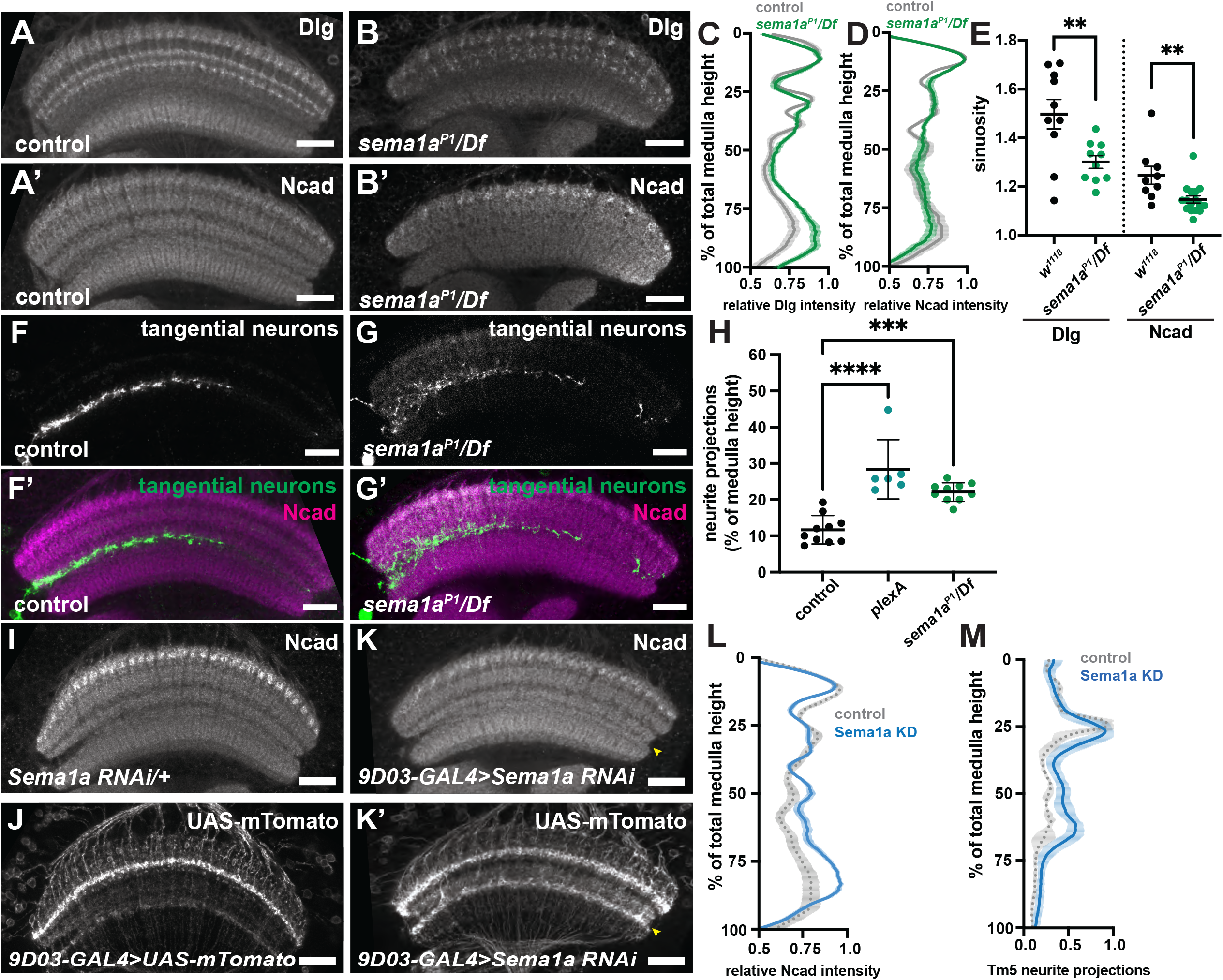
Sema1a is required for layering of the medial medulla. **(A, B)** Control *w^1118^* and *sema1a^P1^*/*Df(2L)Exel7039* mutant 72 h APF brains stained for Dlg **(A, B)** and Ncad **(A’, B’)**. **(C-E)** Sinuosity measurements **(E)** of line intensity traces of Dlg **(C)** and Ncad **(D)** staining in control and *sema1a* mutant medullas. N = 10 control and *sema1a^P1^* mutant Dlg samples; 9 WT, 16 *sema1a^P1^* Ncad samples. **(F, G)** Visualization of tangential neurons with *GMR35A02*-GAL4 driving *UAS-myrTomato* in control **(F)** and *sema1a^P1^*/*Df(2L)Exel7039* **(G)** mutant 72 h APF brains. **(F’, G’)** Tomato shown in green and Ncad in magenta. **(H)** Quantification of the extent of tangential neurites plotted as a percentage of overall medulla height. Measurements were compared via one-way ANOVA with Dunnetts multiple comparisons test, comparing mutant samples to controls. N = 10 WT and *sema1a^P1^*, 6 *plexA^22^* samples. **(I-K)** 72 h APF brains showing *sema1a* knockdown in Tm5 neurons, driven by *GMR9D03*-GAL4 **(K)** compared to *sema1aRNAi/+* **(I)** or *9D03-GAL4/+* **(J)** controls; neuropil labeled with Ncad **(I, K)**, neurites visualized with *UAS*-*myrTomato* **(J, K’)**. Scale bars = 20 µm. **(L)** Quantification of Ncad intensity across medulla height of *sema1a RNAi/+ and 9D03-GAL4>sema1a RNAi* medullas. **(M)** Quantification of Tm5 neurite arborizations in control (*9D03-*GAL4*>UAS-myrTomato*) medullas **(J)** and medullas where *sema1a* is knocked down in Tm5 neurons **(K’)**, plotted as a function of total medulla height. Solid line represents the mean of multiple samples; error bars show SEM. N = 10 *sema1a RNAi/+*, 13 *9D03-GAL4>sema1a RNAi* samples **(L)**; 7 control, 12 *9D03-GAL4>sema1a RNAi* samples **(M)**. **, P ≤ 0.01; ***, P ≤ 0.001; **** P ≤ 0.0001.

While *sema1a^P1^* mutants showed some disruption of medulla layers, they did not fully phenocopy the layering defect observed in *plexA* mutants (compare Fig. 4B with Fig. 3B, E), suggesting that Sema1a could be partially redundant with other Semaphorin family members. We found that both *sema1a, sema1b* and *sema1a; sema5c* double mutants died before pupation, precluding the study of medulla layering in these animals. Therefore, we used a combination of mutant alleles and RNAi knockdown to remove different combinations of Semaphorin proteins from the developing brain. Knockdown of *sema1a* in a *sema1b* mutant, or knockdown of *sema5c* in a *sema1a* mutant did not result in additional layering defects compared to *sema1a* alone (Fig. S3G, H), and *sema1b, sema5c* double mutants did not disrupt medulla layers visualized by Dlg or Ncad staining (Fig. S3F). Thus, we did not find any evidence for redundancy within the Semaphorin family that could explain the differences between *plexA* and *sema1a* mutant layering phenotypes. Recently, a null allele of *sema1a* was generated via CRISPR/Cas9-mediated knockout (*sema1a^SK1^*, National Institute of Genetics Fly Stocks). Surprisingly, analysis of this allele revealed very few homozygous or hemizygous mutant larvae, indicating that complete loss of *sema1a* is lethal, and that the *sema1a^P1^*allele is not null. Examination of medulla layering via Ncad and Dlg staining in an escaper *sema1a^SK1^* hemizygous pupa showed slightly more substantial disruption to layers than was observed in *sema^P1^* mutants, but did not appear as severe as *plexA* (Fig. S3I).

Sema1a has been shown to autonomously control the targeting of lamina neuron projections to the outer medulla by transducing a repulsive activity of PlexA (Pecot et al., 2013). To determine whether the changes in overall layering of the medulla in *sema1a* mutants were a consequence of its function in lamina neurons, we used a lamina-specific driver to knock down expression of *sema1a* and examined the effects on Ncad staining in the medial domain (20-70%) of the medulla. We found only a small effect on overall medulla layering (Fig. S4G, H), although the same RNAi line disrupted layering more severely when expressed in all neurons with two pan-neuronal drivers (Fig. S4A-F), suggesting that loss of Sema1a from medulla neurons contributes to the layering defects. To further identify specific neurons that require Sema1a for layering, we used published single-cell RNA sequencing data (Kurmangaliyev et al., 2020; Ozel et al., 2021) to identify cell types that strongly express *sema1a* during early pupal stages. Of the cells with strong Sema1a expression, we chose to investigate Tm5 neurons because of the availability of an early GAL4 driver. Knockdown of *sema1a* in Tm5 neurons reduced the definition of Ncad layers in the proximal half of the medulla (Fig. 4I, K, L). Further, Tm5 projections were less restricted in layers M5 – 8 than in controls, corresponding to the defects observed in Ncad layering (Fig. 4J, K’, M). Knockdown of *sema1a* in medulla intrinsic Mi1 neurons or transmedullary Tm9 neurons, which have lower levels of *sema1a* expression, did not result in obvious aberrations in medulla layering observed via Ncad staining (Fig. S4I-K); Tm9 arborizations in the distal medulla were largely unaffected (Fig. S4M, O, Q), though Mi1 arborizations in the medial medulla became broader (Fig. S4N-Q). Therefore, Sema1a is required in Tm5 neurons, but not in all medulla neurons, to organize the proximal medulla.

### The cytoplasmic domain of PlexA is not required for lamination of the medulla neuropil

Although PlexA was first described as a receptor for Sema1a, in some contexts their roles are reversed, with PlexA acting as a ligand for Sema1a to activate a downstream “reverse signaling” pathway (Battistini and Tamagnone, 2016).The PlexA cytoplasmic domain is critical for PlexA to function as a receptor; it contains a RasGAP domain and GTPase binding domain (Fig. 5A), and also interacts with other signaling components such as Mical and the guanylyl cyclase Gyc76C (Chak and Kolodkin, 2014; Hung et al., 2010; Yu et al., 2010). In order to determine whether PlexA acts as a receptor in the medulla, we used CRISPR to insert a premature STOP codon 50 amino acids after the transmembrane domain, generating a mutant allele lacking the cytoplasmic domain (*plexA*Δ*cyto)* (Fig. 5A, S5A). An HA tag was added to the C-terminus to allow for monitoring of protein production; Western blotting confirmed the presence of PlexAΔcyto protein in the larval brain (Fig. S5B). HA staining of late-stage pupal brains showed that PlexAΔcyto protein was present in neurites in the medulla, but was not as enriched in the M7 layer as full-length PlexA, suggesting that the cytoplasmic domain contributes to selective trafficking or degradation of the protein (Fig. S5C, D). Surprisingly, *plexA*Δ*cyto* males could survive to adulthood and were fertile, indicating that the receptor function of PlexA is not essential for viability (Fig. S5E).

**Figure 5.**
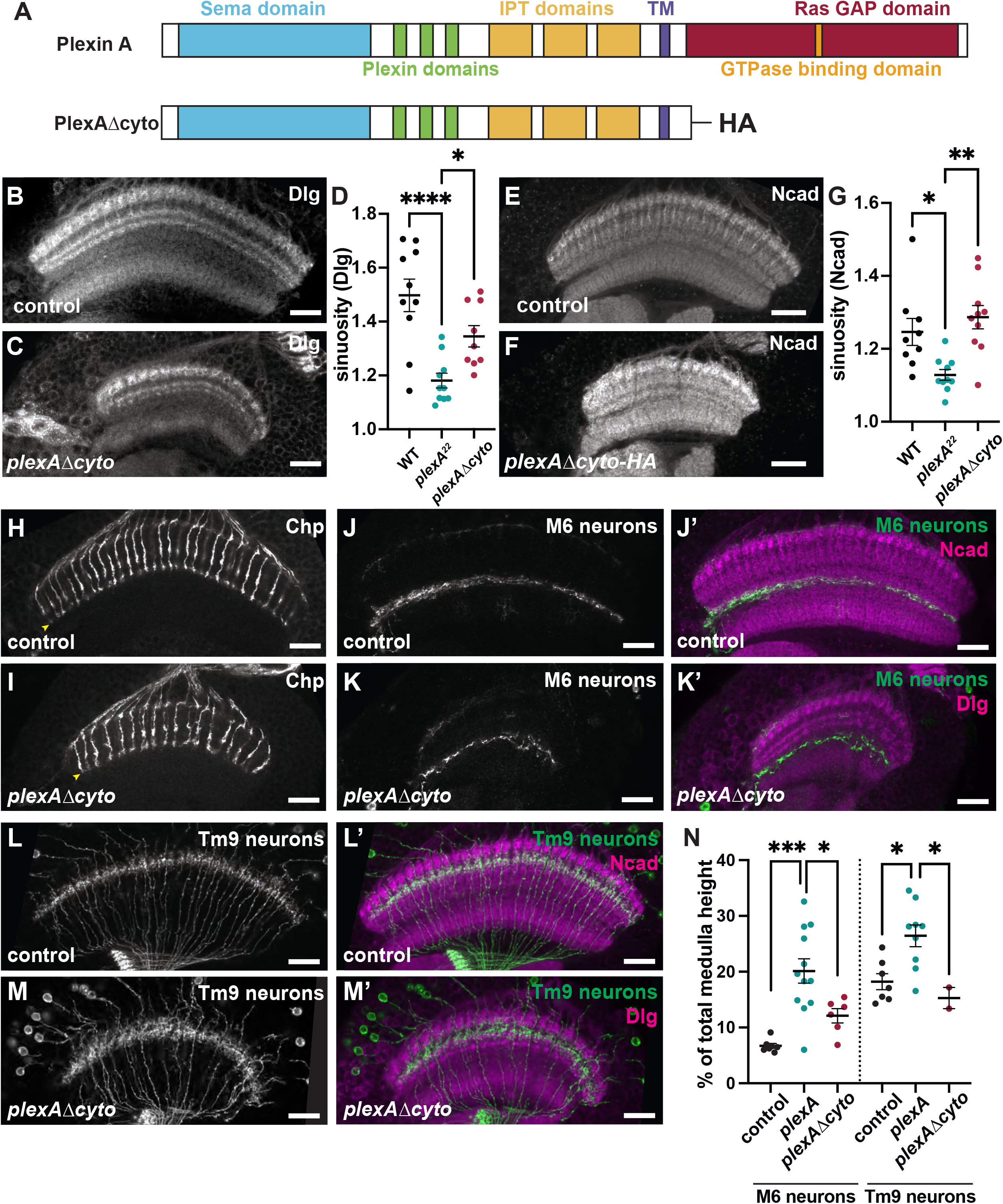
Medulla layers can form in flies lacking the cytoplasmic domain of PlexA. **(A)** Schematic showing the protein domains of full length PlexA and PlexAΔcyto. **(B, E, H, J, L)** show control *w^1118^* and **(C, F, I, K, M)** *plexA*Δ*cyto/plexA^22^* (referred to in this and subsequent figures as *plexA*Δ*cyto*) 72 h APF brains. **(B, C)** Dlg staining; **(E, F)** Ncad staining. **(D, G)** Sinuosity scores of the Dlg **(D)** and Ncad **(G)** intensity curves. N = 10 WT, 10 *plexA^22^*, 9 *plexA*Δ*cyto* Dlg samples; 9 WT, 10 *plexA^22^*, 10 *plexA*Δ*cyto* Ncad samples. **(H, I)** R7/8 photoreceptor axons visualized by Chp staining in *w^1118^* control **(H)** and *plexA*Δ*cyto* mutant brains **(I)**. **(J, K)** M6 local medulla neurons visualized by *30A-*GAL4 and **(L, M)** Tm9 neurons visualized by GMR*24C08-*GAL4 driving *UAS-myrTomato*. In **(J’, K’, L’, M’)** Tomato is shown in green and the neuropil is labeled with Ncad **(J’, L’)** or Dlg **(K’, M’)** in magenta. Scale bar = 20 µm. **(N)** Quantification of the extent of M6 and Tm9 neurite outgrowth in *w^1118^*, *plexA^22^/plexA^MB09499^* and *plexA*Δ*cyto* mutants. N = 7 control, 12 *plexA*, 6 *plexA*Δ*cyto* M6 samples; 7 WT, 9 *plexA*, 2 *plexA*Δ*cyto* Tm9 samples. Measurements were compared via one-way ANOVA with Tukey’s multiple comparisons test, comparing the means of each group. *, P ≤ 0.05; **, P ≤ 0.01; ***, P ≤ 0.001; if no significance is reported, differences between groups were not significant.

To test the role of the cytoplasmic domain of PlexA in medulla layering, we stained *plexA*Δ*cyto* pupal brains for Dlg, Ncad and Sdk, as well as Connectin (Con), a homophilic adhesion protein involved in axon guidance, and Choline acetyltransferase (ChAT) to reveal the pattern of layers (Fig. 5B-G, Fig. S5C’, D’, F-J). Surprisingly, the layers observed with these markers were more distinct than in *plexA* null mutants (Fig. 5D, G; Fig. S5H, K). In addition, R7 photoreceptors terminated correctly in the M6 layer (Fig. 5H, I), and M6 and Tm9 neurons showed almost normal arborization patterns in *plexA*Δ*cyto* mutants (Fig. 5J-N), in contrast to the disordered projections observed in *plexA* null mutants (Fig. 5N). These data show that PlexA expressed at endogenous levels can mediate substantial layering without its cytoplasmic domain, making it likely that it acts as a ligand rather than a receptor in this context. Tangential neuron projections remained significantly disordered in *plexA*Δ*cyto* mutant pupae (Fig. S5L-N), suggesting that tangential neuron targeting requires an autonomous receptor function of PlexA.

### PlexA is required for normal morphology of the optic lobe neuropils

In addition to establishing stereotyped synaptic layers, the medulla neuropil undergoes substantial morphological changes during pupal development. In early pupal stages, transverse optical sections of the medulla neuropil stained for Ncad have a distinct wedge shape, where the narrow part of the wedge corresponds to the nascent posterior neuropil, and the wider anterior edge is further developed (Ngo, Andrade and Hartenstein, 2017)(Fig. 6A). As development proceeds, the medulla loses its wedge shape and acquires an even height throughout when viewed in horizontal sections (Fig. 6B). After 48 h APF, the general shape of the medulla remains constant, but it increases in height as synaptic layers increase in resolution (Ngo, Andrade and Hartenstein, 2017) (Fig. 6C).

**Figure 6.**
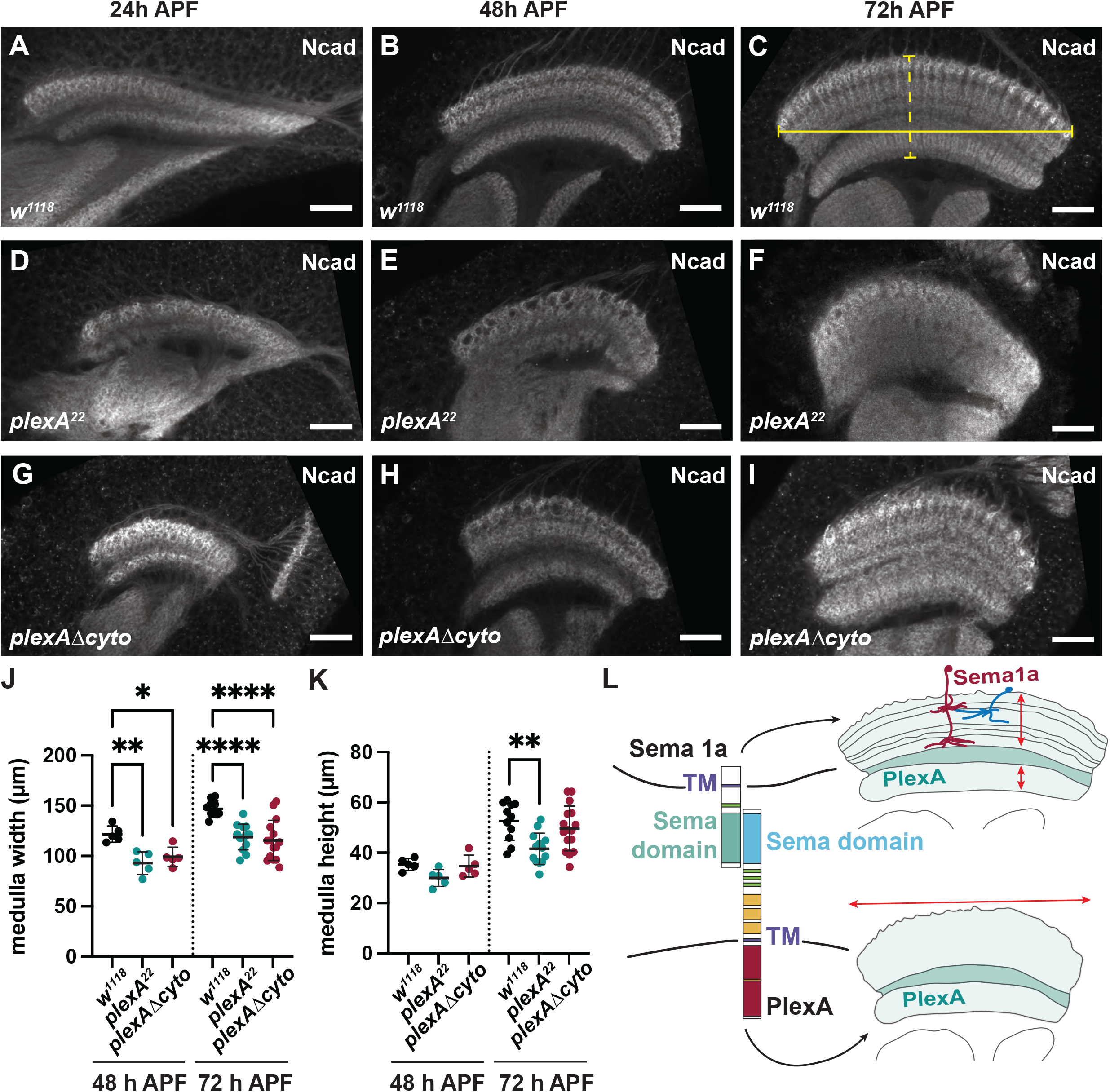
The cytoplasmic domain of PlexA is required for normal medulla morphogenesis. Whole-mounted pupal brains stained for Ncad to visualize the medulla neuropil along a horizontal plane at 24 h APF **(A, D, G)**, 48 h APF **(B, E, H)**, and 72 h APF **(C, F, I)**. **(A-C)** *w^1118^* control; **(D-F)** *plexA^22^*; **(G-I)** *plexA*Δ*cyto*. Scale bar = 20 µm. **(J)** Quantification of medulla anterior-posterior width (solid yellow bracket in **C**) at 48 h APF and 72 h APF in control, *plexA^22^*, and *plexA*Δ*cyto* mutants. **(K)** Quantification of medulla distal-proximal height (dashed yellow bracket in **C**) at 48 h APF and 72 h APF in control, *plexA^22^*, and *plexA*Δ*cyto* mutant pupal brains. Measurements were compared via one-way ANOVA with Dunnetts multiple comparisons test, comparing mutant samples to wild-type controls. N = 5 48 h APF samples/ genotype; 11 control, 13 *plexA^22^*, 15 *plexA*Δ*cyto* 72 h APF samples. Error bars denote mean ± SD. *, P ≤ 0.05; **, P ≤ 0.01; ****, P ≤ 0.0001. **(L)** Proposed model for a dual role for PlexA in regulating lamination and morphogenesis of the medulla neuropil. PlexA acts as a ligand for Sema1a expressed in a subset of neurons (red) such as Tm5 that then act as a scaffold to guide the projections of other neurons (blue). PlexA also acts as a receptor to control the overall shape of the medulla neuropil.

To characterize how loss of *plexA* affects medulla morphogenesis, we stained early-, mid-, and late-stage pupal brains for Ncad to label the neuropil and quantified the anterior-posterior width and proximal-distal height of the medulla neuropil in horizontal sections (Fig. 6A-K). Medullas lacking all *plexA* displayed a notably less pronounced wedge shape at 24 h APF than controls (Fig. 6A, D), suggesting that *plexA* is required as early as the onset of pupal development to establish the correct neuropil shape. Pupal medullas homozygous for *plexA^22^* showed significant decreases in both width and height at 72 h APF, and in width at 48 h APF, compared to controls (Fig. 6B, C, E, F, J, K). The *plexA^MB09499^* allele similarly altered the shape of the medulla neuropil (Fig. S6A, B, E, F). To determine whether the total size of the neuropil was reduced, we analyzed frontal sections of 72 h APF pupal brains. In control samples, the medulla is located distal to the lobula, forming an arc above the lobula and lobula plate neuropils (Fig. S6G, J). In *plexA* mutants the medulla increased in length along the dorsal-ventral axis, wrapping around the lobula complex in a semicircular shape (Fig. S6H, J). This suggests that the defect in *plexA* mutants is a change in the aspect ratio of the medulla neuropil rather than a reduction in its volume.

We next asked whether PlexA regulates medulla morphogenesis as a ligand or a receptor. While loss of the PlexA cytoplasmic domain did not significantly affect the height of the medulla neuropil, consistent with its limited effect on the segregation of medulla layers, the width of horizontal sections of the medulla neuropil in *plexA*Δ*cyto* pupal brains was reduced to a similar extent as in *plexA* null mutants (Fig. 6G-K). Examination of *plexA*Δ*cyto* mutants in frontal cross-sections revealed an increase in medulla length along the dorsal-ventral axis (Fig. S6G, J), similar to *plexA* null mutants. As observed in *plexA^22^* mutants, the medulla morphology defects in *plexA*Δ*cyto* mutants were present as early as 24 h APF and persisted throughout mid- and late-pupal development (Fig. 6G-I), suggesting that medulla morphogenesis requires PlexA to act as a receptor. The decrease in medulla width seen in *plexA*Δ*cyto* and *plexA^22^* mutants was not observed to the same extent in *sema1a^P1^* mutants (Fig. S6C-F), indicating that Sema1a may not be the only ligand for this function of PlexA. Thus, PlexA likely acts as a ligand for Sema1a to segregate synaptic layers in the medulla, but as a receptor, perhaps for a different ligand, to give the medulla neuropil its overall shape during pupal development (Fig. 6L).

## Discussion

Here we describe two distinct roles for PlexA in optic lobe development. In a null *plexA* mutant, the medulla fails to form its normal pattern of ten distinct synaptic layers. Loss of *sema1a* has a similar effect, suggesting that PlexA acts at least in part through Sema1a to mediate medulla lamination. Importantly, the cytoplasmic domain of PlexA is dispensable for establishing medulla layering, indicating that PlexA acts as a ligand rather than a receptor in this context. In contrast, the cytoplasmic domain of PlexA is required to define the overall shape of the medulla neuropil.

### PlexA has separate ligand and receptor functions

In addition to their well-defined role as ligands for Plexins, transmembrane Semaphorins have been implicated as receptors that activate a reverse signaling pathway in several vertebrate and invertebrate systems (Battistini and Tamagnone, 2016) (Andermatt et al., 2014; Sun et al., 2015; Toyofuku et al., 2004; Yu et al., 2010). By removing the cytoplasmic domain from endogenous PlexA protein via CRISPR mutagenesis, we provide conclusive evidence that this domain is not essential for medulla layer formation *in vivo*, consistent with PlexA acting as a ligand for Sema1a in this process (Fig. 6L).

In contrast, the cytoplasmic domain of PlexA is required for normal morphogenesis of the medulla neuropil, suggesting that this process is mediated by a more canonical receptor function of PlexA. Defects in medulla size and shape are observed in early pupal stages in *plexA* mutants, indicating that PlexA acts during the initial establishment of the neuropil structure. The cells in which PlexA acts to define the neuropil boundary are unknown. In the fly antennal lobe, discrete compartments are established by ensheathing glia that wrap each neuropil compartment in response to FGF signaling (Wu et al., 2017). Loss of this glial barrier disrupts neuronal targeting to the correct region of the antennal lobe (Wu et al., 2017), raising the possibility that the establishment of the correct medulla neuropil shape also requires crosstalk between glia at the boundary and incoming neuronal processes.

The ligand for PlexA in the control of medulla morphogenesis remains to be identified. Loss of Sema1a, the ligand used in many other contexts (Winberg et al., 1998; Winberg et al., 2001; Yu et al., 1998), does not alter medulla neuropil shape to the same extent as loss of PlexA. It is possible that Sema1a acts redundantly with other Semaphorins to direct medulla morphogenesis; the early lethality of combinations of *sema* mutations prevented us from testing this hypothesis. Another possible ligand is the secreted protein Slit, which is required in medulla neurons and glia to regulate morphogenesis of the optic lobe neuropils and establish boundaries between them (Caipo et al., 2020; Suzuki et al., 2016; Tayler et al., 2004). At the mouse midline, a C-terminal fragment of Slit has been shown to bind to PlexinA1, inducing repulsive activity that is independent of canonical Robo or Neuropilin signaling pathways (Delloye-Bourgeois et al., 2015).

Surprisingly, homozygous mutants lacking the PlexA cytoplasmic domain are viable. Thus, while PlexA acts as a receptor in many processes in *Drosophila* including motor axon guidance (Winberg et al., 1998), development of olfactory circuits (Sweeney et al., 2007), cell migration in the ovary (Stedden et al., 2019), and epithelial repair after wound healing (Yoo et al., 2016), its receptor function is not critical for survival.

### PlexA signaling through Sema1a refines synaptic layers in the medulla

PlexA was previously reported to restrict lamina neuron projections to the distal domain of the medulla (Pecot et al., 2013), based on the effects of *plexA* RNAi. Our results extend this analysis to other medulla neurons, and examine the relative effects of PlexA protein domains on the lamination and shape of the entire medulla neuropil. We found that loss of *plexA* affects not only lamina neurons, but many other neurites that project into the medulla, leading to broader and overlapping synaptic layers. This is similar to the effect of *plexA* on the ellipsoid body, a neuropil substructure located in the central brain that is innervated by ring neurons (Hanesch et al., 1989). Knockdown of either *sema1a* or *plexA* in subsets of ring neurons disrupts the strict segregation of laminae in the ellipsoid body (Xie et al., 2017). Sema1a appears to act as a receptor in R4m ring neurons to prevent them from forming synapses in the central ellipsoid body, and loss of *sema1a* from these neurons in turn affects R3 neurons, which expand their arborizations into the outer ellipsoid body (Xie et al., 2017). A similar non-autonomous function might explain the patterning of the medulla by PlexA and Sema1a. Single cell RNA sequencing data indicate that although *sema1a* is expressed in many different neuronal subtypes throughout the medulla (Kurmangaliyev et al., 2020; Ozel et al., 2021), it is not uniformly expressed in all cells. Further, we show that removal of *sema1a* from Tm5 neurons is sufficient to disrupt regional medulla layering, while other cell types such as Tm9 neurons do not autonomously require *sema1a*. It is possible that Tm5 neurons and other specific Sema1a-expressing cells respond directly to the PlexA ligand, and that their projections establish a scaffold that indirectly affects the lamination of other neuronal processes (Fig. 6L).

While some molecular cues that establish synaptic lamination are themselves segregated into layers (Yamagata and Sanes, 2008; Yamagata and Sanes, 2012), this does not seem to be the case for PlexA. PlexA protein is enriched in the M7 layer and its effects on layer formation appear weakest in the layers furthest from M7, but it is also present at lower levels throughout the medulla. Our somatic CRISPR experiments support a requirement for PlexA in tangential neuron precursors, but do not exclude a role for it in other neurons, as the knockout achieved by this method is unlikely to be complete. The widespread distribution of PlexA seems to exclude a model in which PlexA is always a repulsive signal, as in this case Sema1a-expressing neurites would fail to enter the medulla neuropil. It is possible that different levels of PlexA have distinct attractive or repulsive effects, or that PlexA has a function *in cis* in Sema1a-expressing cells (Rozbesky et al., 2020; Sun et al., 2013). PlexA and Sema1a could also influence one another’s location through mechanisms such as localized binding or receptor-mediated endocytosis, leading to local differences in protein levels that might not be detectable by antibody staining but could contribute to the establishment of layer boundaries. Additionally, the dynamic nature of medulla layering (Ngo, Andrade and Hartenstein, 2017) means that a growing neurite may receive locational cues from cells that later change their position or gene expression, rendering the temporal as well as spatial patterns of protein distribution relevant for targeting. Interestingly, loss of the PlexA cytoplasmic domain results in dispersed tangential neuron projections, suggesting a cell-autonomous role for PlexA as a receptor in the bundling of tangential neuron processes that might explain its higher expression in the M7 layer.

A variety of roles for Plexin-Semaphorin signaling in layer-specific neurite arborization in the mouse retina and in retinal ganglion cell targeting have been described (Matsuoka et al., 2011; Sun et al., 2015; Sun et al., 2013). Our findings provide further evidence that interactions between Plexins and Semaphorins can regulate neuropil organization via both cell autonomous and non-autonomous mechanisms, depending on the context, and strengthen the findings of structural and functional homology between the fly optic lobe and mouse retina (Sanes and Zipursky, 2010). Further, the role of PlexA in medulla neuropil morphogenesis establishes a molecular connection between the gross morphology of a brain structure and its internal synaptic organization.

## Materials and Methods

### Fly stocks and genetics

Fly stocks used were *plexA^22^ (this paper)*; *plexA^MB09499^ (BDSC#61741); plexAΔcyto-HA* (this paper); *plexA*Δ*Sema* (this paper); *sema1a^P1^*(BDSC#11097); *sema1a^SK1^* (NIG-Fly#M2L-3127) *sema1b^KO^*(Wittes and Schüpbach, 2019); *sema5c^K175^* (Stedden et al., 2019); *Df(2L)Exel7039* (BDSC#7810) *UAS-CD8-GFP* (Kyoto #108068); *UAS-myrTomato* (BDSC#32221); *UAS-dcr2* (BDSC#24650); *UAS-Cas9-P2* (from BDSC#54593); *repo-GAL4* (BDSC#7415); *nSyb-GAL4* (BDSC#51635); *elav-GAL4* (BDSC #8765); *hth-GAL4* (Wernet et al., 2003); *Dll-GAL4* (BDSC #3038); *30A-GAL4* (BDSC#37534); *GMR35A02-GAL4* (BDSC#49811); *GMR24C08-GAL4* (BDSC#48050); *ort^C2b^-GAL4* (Ting et al., 2014); *GH146-GAL4* (BDSC#30026); *27G05-FLP, Act>CD2>GAL4* (Pecot et al., 2013); *GMR9B08-GAL4* (BDSC#41369); *Rh3-LexA* (gift of C. Desplan); *LexAop::brpShort-mCherry* (Mosca and Luo, 2014); *sema1a RNAi* HMS01307 (BDSC#34320); *sema5c RNAi* JF03372 (BDSC#29436); *plexA RNAi* HM05221 (BDSC#30482); *plexA RNAi* KK101499 (VDRC#v107004); *pCFD4-plexA sgRNAs* (this paper); *UAS-plexA-HA* (Terman et al., 2002); *UAS-plexAΔcyto-HA* (Hung et al., 2010); *w^1118^* (BDSC#3605); *Bac-PlexA-Myc* (Pecot et al., 2013).

The genotype used to generate large *plexA* mutant clones in a *Minute* background was *ey3.5-FLP; tub-GAL4, UAS-myrTomato; FRT80, Rh3/4-lacZ / FRT80, M(3)67C, tub-GAL80, Dp(4;3)RC051; plexA^MB09499^*.

*repo-GAL4* was used to drive gene expression in glial cells; *elav-GAL4* and *nSyb-GAL4* were used to drive gene expression in neurons. *hth-GAL4 and Dll-GAL4* were used to drive gene expression in medulla and tangential neuron precursors, respectively (Bertet et al., 2014). *GMR35A02-GAL4* was used to label tangential neurons in the pupal medulla. *30A-GAL4* was used to label M6 local medulla neurons in the pupa (Chin et al., 2014). *GMR24C08-GAL4* was used to label Tm9 neurons (Chin et al., 2014). Dm8 neurons were labeled with *ort^C2b^-GAL4* (Ting et al., 2014). *GMR9D03-GAL4* was used to label Tm5 neurons (Han et al., 2011). *bshM-GAL4* was used to label Mi1 neurons (Trush et al., 2019). *GMR9B08-GAL4* was used to drive gene expression in lamina neurons (Pecot et al., 2013; Schwabe et al., 2014). *w^1118^*flies were used as a wildtype control throughout.

### Cloning and transgenic lines

To generate the *plexA^22^* mutant allele, the *plexA* sgRNA sequences CATTACTTCAGTACCGGTGG and GTTGACGCTTGTACACATGA were made by gene synthesis in pUC57 (GenScript) and cloned into pCFD4 (Port et al., 2014) by Gibson assembly. The pCFD4-null construct was integrated into the attP40 site at 25C6. These flies were crossed to *nos-Cas9* flies to make germline mosaic flies. The progeny of these flies were crossed to balancer flies and screened first for failure to complement *plexA^MB09499^*, then by PCR for the expected deletion, using the primers GGCGTGGCTATTCGTTATTCG and TGCGACACAATTGGACAAGTG.

To generate the *plexA*Δ*Sema* mutant, the sema sgRNA sequences ACTTTCCTTATTGTTTTTTC and TGTGTTCTAACGGTAAGTAT were made by gene synthesis in pUC57 (GenScript) then cloned into pCFD4 (Port et al., 2014) by Gibson assembly. The pCFD4-Sema construct was integrated into the attP40 site at 25C6. Injections and screening of transgenic flies were carried out by GenetiVision. After crossing to *nos-Cas9* flies and screening by PCR, the deletion expected if the DNA had been cut at both sgRNA positions was not obtained. The lethal mutations were characterized by PCR using the primers CGTAGCATTTATTGCGCCG and CGTGGGAATTAACTTTAAAGGGC. We isolated one mutation that had a single base-pair insertion and one that had a two base-pair deletion just before the 3’ splice junction of the Sema domain-encoding exon. Both introduced a frameshift leading to a stop codon shortly thereafter, and would prevent transcripts containing this exon from producing a functional protein. Fig. S3A, B show the single base-pair deletion.

The *plexA*Δ*cyto* mutant was generated via CRISPR mutagenesis by GenetiVision Corporation. A pCFD5 plasmid containing gRNA1 (Kanca et al., 2019) and a gRNA that targeted the *PlexA* gene just after the transmembrane domain (CTTCGCTCACAGGCGATCTTACTTC) was injected into *nos-Cas9* expressing flies together with a pUC57 plasmid containing a donor construct (synthesized by GENEWIZ/Azenta) flanked by gRNA1 target sites that would insert an HA tag and 3X STOP cassette. Successful insertion of the donor fragment was confirmed by PCR amplification and sequencing.

### Histology

Pupal development stages were calculated from the white prepupal stage (0 h APF) at 25°C. Pupal brains were dissected in cold PBS, fixed in 4% paraformaldehyde in PBS for 30 min at room temperature, and washed 3 times for 5 min in PBST (PBS + 0.3% TritonX-100). Adult brains were dissected for whole mount in cold Schneider’s medium, and fixed in 4% paraformaldehyde in PBS for 30 minutes at room temperature. After washing, samples were blocked in PBST + 10% donkey serum for 1 h prior to incubation with primary antibodies in PBST overnight (pupal brains) or for 72 h (adult brains) at 4°C. Samples were washed in PBST three times for 20 min each and incubated in secondary antibodies for 4 h at room temperature, or overnight at 4°C. Samples were washed in PBST three times for 20 min each and mounted in SlowFade Gold AntiFade reagent (Invitrogen) on bridge slides. Confocal images were collected on a Leica SP8 confocal microscope using a 63x/1.40 oil objective. Images were processed in FIJI and Adobe Photoshop.

Adult heads were dissected for cryosection in cold 0.1 M sodium phosphate buffer (PB) pH 7.4, fixed in 4% formaldehyde in PB for 4 h at 4°C and washed in PB. Heads were then submerged in a sucrose gradient (5%, 10%, 20%) for 20 min at each concentration, and left in 25% sucrose overnight at 4°C for cryoprotection. Heads were embedded in OCT tissue freezing medium and frozen in dry ice/ethanol, and 12 μm sections were cut on a cryostat. Sections were post-fixed in 0.5% formaldehyde in PB for 30 min at room temperature and washed three times in PB with 0.1% Triton (PBT) before incubation with primary antibodies overnight at 4°C. Sections were washed four times for 20 min with PBT and incubated with secondary antibodies for 2 h at room temperature. Sections were washed again four times for 20 min before mounting in Fluoromount-G (Southern Biotech).

Primary antibodies used were mouse anti-Chp (1:50, Developmental Studies Hybridoma Bank [DSHB] 24B10), chicken anti-GFP (1:400, Invitrogen), rat anti-HA (1:50, Roche 3F10), mouse anti-HA (1:200; BioLegend), rat anti-Ncad (1:50, DSHB Ex#8), rabbit anti-dsRed (1:500, Takara Bio), guinea pig anti-Sdk (1:200) (Astigarraga et al., 2018), mouse anti-ChAT (1:50, DSHB 4B1), mouse anti-Con (1:50, DSHB C1.427), mouse anti-Dlg (1:20, DSHB 4F3), rabbit anti-PlexA (1:500, gift of L. Luo) (Sweeney et al., 2007), or rabbit anti-Myc (1:1000, Abcam). Secondary antibodies used were coupled to AlexaFluor488 (Invitrogen), Cy3 (Jackson Immunoresearch), Cy5 (Jackson Immunoresearch), or AlexaFluor647 (Invitrogen).

### Western blotting

To extract proteins, L3 larval brains were dissected and homogenized in lysis buffer [50 mM Tris-HCl (pH 8), 150 mM NaCl, 1% Triton X-100, 0.5% deoxycholate, 0.1% SDS, 5 mM NaF, 1mM Na3VO4 (pH 9.96), 5 mM EDTA, Complete Protease Inhibitor Cocktail (Roche)]. Samples were mixed with 4X Laemmli buffer [4% SDS, 20% glycerol, 120 mM Tris-Cl pH 6.8, 0.02% bromophenol blue, 10% beta-mercaptoethanol] and heated at 95°C for 3 min before loading onto an SDS-PAGE gel. Gels were run first at 85 volts for 15 min, then 150 volts for the remainder of the time and transferred onto 0.45 µm nitrocellulose membranes (BioRad) for 2.5 hours at 75 volts. Membranes were washed for 5 min in TBS and blocked in 5% low-fat milk in TBST (20 mM Tris (pH 7.6), 136 mM NaCl, 0.2% Tween-20) solution for 1 h. Membranes were incubated overnight with primary antibody in TBST with 5% milk at 4°C, washed three times for 10 min in TBST, and incubated in horseradish peroxidase-conjugated secondary antibodies (1:2,000, Jackson ImmunoResearch) at room temperature in TBST with 5% milk for 2 h. Membranes were washed three times for 10 min in TBST and once for 10 min in TBS. Blots were developed with enhanced chemiluminescence (Thermo SuperSignal WestPico). Primary antibodies used were rat anti-Elav (1:2,000, DSHB 7E8A10), rat anti-HA (1:2,000, Roche 3F10), mouse anti-β-tubulin (1:10,000; Sigma T4026), and rabbit anti-PlexA (1:2,000, (Sweeney et al., 2007)).

### Quantification of images

All quantifications were done on maximum projections of 10 confocal sections (3 µm thick total stack). To quantify medulla layering in brains stained for Ncad, Dlg, Sdk, ChAT, and Con, and neurite outgrowth in medulla and tangential neurons, the Straighten function in ImageJ was used to straighten a region of the medulla representing greater than 50% of the overall neuropil. Image intensity of neuropil markers or neurite projections was measured over that region of interest using Plot Profile in FIJI, and the data array was resampled to include 300 data points, normalizing the medulla height of each sample. Intensity values were normalized by dividing all values in a sample by the highest value in that sample. Mean intensity profiles were plotted as a function of relative medulla height. The % neurite outgrowth was calculated by measuring the width of the corresponding intensity peak at 50% height, and dividing by the medulla height; measurements of each condition were averaged together.

Measurements of overall medulla morphology were taken in FIJI, by measuring the length of a user-defined line from the distal-most anterior to posterior corners (medulla width), and distal to proximal extent at the center-most position (medulla height). Area measurements were obtained by measuring the area of a user-defined polygon in FIJI.

## Acknowledgements

We thank Claude Desplan, Isabel Holguera, Sally Horne-Badovinac, Alex Kolodkin, Chi-Hon Lee, Liqun Luo, Makoto Sato, Takashi Suzuki, Julia Wittes, Larry Zipursky, the Bloomington *Drosophila* Stock Center, the Vienna *Drosophila* Resource Center, the Kyoto Stock Center, the National Institute for Genetics, and the Developmental Studies Hybridoma Bank for fly stocks and reagents, and Flybase for essential information. We are grateful to DanQing He, Ariel Hairston, and Dhaval Gandhi for expert technical assistance, to Michael Cammer for helpful discussions and assistance with image analysis, and to the NYU Microscopy Core for technical support. The NYU Microscopy Core (RRID: SCR_01934) is partially supported by the Cancer Center Support Grant P30CA016087 at the Laura and Isaac Perlmutter Cancer Center; instruments supported by NIH grant S10 RR023708. The manuscript was improved by the critical comments of Neha Ghosh, Hongsu Wang, Chris Doe, Katherine Nagel, and Niels Ringstad. This work was supported by the National Institutes of Health (grant R01 NS112211 to J.E.T. with supplement NS112211-02S1 to M.E.B., and fellowship F31 EY025568 to J.D.).

## References

Andermatt, I., Wilson, N. H., Bergmann, T., Mauti, O., Gesemann, M., Sockanathan, S. and Stoeckli, E. T. (2014) Semaphorin 6B acts as a receptor in post-crossing commissural axon guidance. Development, 141, 3709–3720.

Astigarraga, S., Douthit, J., Tarnogorska, D., Creamer, M. S., Mano, O., Clark, D. A., Meinertzhagen, I. A. and Treisman, J. E. (2018) *Drosophila* Sidekick is required in developing photoreceptors to enable visual motion detection. Development, 145.

Astigarraga, S., Hofmeyer, K. and Treisman, J. E. (2010) Missed connections: photoreceptor axon seeks target neuron for synaptogenesis. Curr Opin Genet Dev, 20, 400–407.

Ayoob, J. C., Yu, H. H., Terman, J. R. and Kolodkin, A. L. (2004) The *Drosophila* receptor guanylyl cyclase Gyc76C is required for Semaphorin-1a-Plexin A-mediated axonal repulsion. J. Neurosci., 24, 6639–6649.

Baier, H. (2013) Synaptic laminae in the visual system: molecular mechanisms forming layers of perception. Annu Rev Cell Dev Bi, 29, 385–416.

Battistini, C. and Tamagnone, L. (2016) Transmembrane Semaphorins, forward and reverse signaling: have a look both ways. Cell Mol Life Sci, 73, 1609–1622.

Bellen, H. J., Levis, R. W., He, Y., Carlson, J. W., Evans-Holm, M., Bae, E., Kim, J., Metaxakis, A., Savakis, C., Schulze, K. L., et al (2011) The *Drosophila* gene disruption project: progress using transposons with distinctive site specificities. Genetics, 188, 731–743.

Bertet, C., Li, X., Erclik, T., Cavey, M., Wells, B. and Desplan, C. (2014) Temporal patterning of neuroblasts controls Notch-mediated cell survival through regulation of Hid or Reaper. Cell, 158, 1173–1186.

Caipo, L., Gonzalez-Ramirez, M. C., Guzman-Palma, P., Contreras, E. G., Palominos, T., Fuenzalida-Uribe, N., Hassan, B. A., Campusano, J. M., Sierralta, J. and Oliva, C. (2020) Slit neuronal secretion coordinates optic lobe morphogenesis in *Drosophil*a. Dev Biol, 458, 32–42.

Chak, K. and Kolodkin, A. L. (2014) Function of the *Drosophila* receptor guanylyl cyclase Gyc76C in PlexA-mediated motor axon guidance. Development, 141, 136–147.

Chin, A. L., Lin, C. Y., Fu, T. F., Dickson, B. J. and Chiang, A. S. (2014) Diversity and wiring variability of visual local neurons in the *Drosophila* medulla M6 stratum. J. Comp. Neurol., 522, 3795–3816.

Fischbach, K. F. and Dittrich, A. P. M. (1989) The optic lobe of *Drosophila melanogaster*. I. A Golgi analysis of wild-type structure. Cell Tissue Res., 258.

Fisher, Y. E., Leong, J. C., Sporar, K., Ketkar, M. D., Gohl, D. M., Clandinin, T. R. and Silies, M. (2015) A class of visual neurons with wide-field properties is required for local motion detection. Curr Biol, 25, 3178–3189.

Gao, S., Takemura, S.-y., Ting, C.-Y., Huang, S., Lu, Z., Luan, H., Rister, J., Thum, A. S., Yang, M., Hong, S.-T., et al (2008) The neural substrate of spectral preference in *Drosophila*. Neuron, 60, 328–342.

Guy, J. and Staiger, J. F. (2017) The functioning of a cortex without layers. Front Neuroanat, 11, 54.

Han, C., Jan, L. Y. and Jan, Y. N. (2011) Enhancer-driven membrane markers for analysis of nonautonomous mechanisms reveal neuron-glia interactions in *Drosophila*. Proc Natl Acad Sci U S A, 108, 9673–9678.

Hanesch, U., Fischbach, K. F. and Heisenberg, M. (1989) Neuronal architecture of the central complex in *Drosophila melanogaster*. Cell Tissue Res., 257, 343–366.

Hung, R.-J., Yazdani, U., Yoon, J., Wu, H., Yang, T., Gupta, N., Huang, Z., van Berkel, W. J. H. and Terman, J. R. (2010) Mical links semaphorins to F-actin disassembly. Nature, 463, 823–827.

Janssen, B. J. C., Robinson, R. A., Pérez-Brangulí, F., Bell, C. H., Mitchell, K. J., Siebold, C. and Jones, E. Y. (2010) Structural basis of Semaphorin–Plexin signalling. Nature, 467, 1118–1122.

Jenett, A., Rubin, G. M., Ngo, T. T., Shepherd, D., Murphy, C., Dionne, H., Pfeiffer, B. D., Cavallaro, A., Hall, D., Jeter, J., et al (2012) A GAL4-driver line resource for *Drosophila* neurobiology. Cell Rep, 2, 991–1001.

Jeong, S., Juhaszova, K. and Kolodkin, Alex L. (2012) The control of Semaphorin-1a-mediated reverse signaling by opposing Pebble and RhoGAPp190 functions in *Drosophila*. Neuron, 76, 721–734.

Jeong, S., Yang, D. S., Hong, Y. G., Mitchell, S. P., Brown, M. P. and Kolodkin, A. L. (2017) Varicose and Cheerio collaborate with Pebble to mediate Semaphorin-1a reverse signaling in *Drosophila*. Proc Natl Acad Sci U S A, 114, E8254–E8263.

Kanca, O., Zirin, J., Garcia-Marques, J., Knight, S. M., Yang-Zhou, D., Amador, G., Chung, H., Zuo, Z., Ma, L., He, Y., et al (2019) An efficient CRISPR-based strategy to insert small and large fragments of DNA using short homology arms. Elife, 8, e51539.

Kind, E., Longden, K. D., Nern, A., Zhao, A., Sancer, G., Flynn, M. A., Laughland, C. W., Gezahegn, B., Ludwig, H. D. F., Thomson, A. G., et al (2021) Synaptic targets of photoreceptors specialized to detect color and skylight polarization in *Drosophila*. eLife, 10, e71858.

Kurmangaliyev, Y. Z., Yoo, J., Valdes-Aleman, J., Sanfilippo, P. and Zipursky, S. L. (2020) Transcriptional programs of circuit assembly in the *Drosophila* visual system. Neuron, 108, 1045–1057.

Kurusu, M., Cording, A., Taniguchi, M., Menon, K., Suzuki, E. and Zinn, K. (2008) A screen of cell-surface molecules identifies leucine-rich repeat proteins as key mediators of synaptic target selection. Neuron, 59, 972–985.

Li, X., Erclik, T., Bertet, C., Chen, Z., Voutev, R., Venkatesh, S., Morante, J., Celik, A. and Desplan, C. (2013) Temporal patterning of *Drosophila* medulla neuroblasts controls neural fates. Nature, 498, 456–462.

Liu, H., Juo, Z. S., Shim, A. H., Focia, P. J., Chen, X., Garcia, K. C. and He, X. (2010) Structural basis of Semaphorin-Plexin recognition and viral mimicry from Sema7A and A39R complexes with PlexinC1. Cell, 142, 749–761.

Matsuoka, R. L., Nguyen-Ba-Charvet, K. T., Parray, A., Badea, T. C., Chédotal, A. and Kolodkin, A. L. (2011) Transmembrane Semaphorin signalling controls laminar stratification in the mammalian retina. Nature, 470, 259–263.

Menon, K. P., Kulkarni, V., Takemura, S. Y., Anaya, M. and Zinn, K. (2019) Interactions between Dpr11 and DIP-gamma control selection of amacrine neurons in *Drosophila* color vision circuits. Elife, 8, e48935.

Millard, S. S. and Pecot, M. Y. (2018) Strategies for assembling columns and layers in the *Drosophila* visual system. Neural Dev, 13, 11.

Morante, J. and Desplan, C. (2008) The color-vision circuit in the medulla of *Drosophila*. Current Biology, 18, 553–565.

Mosca, T. J. and Luo, L. (2014) Synaptic organization of the *Drosophila* antennal lobe and its regulation by the Teneurins. Elife, 3, e03726.

Negishi, M., Oinuma, I. and Katoh, H. (2005) Plexins: axon guidance and signal transduction. Cell. Mol. Life Sci., 62, 1363.

Ngo, K. T., Andrade, I. and Hartenstein, V. (2017) Spatio-temporal pattern of neuronal differentiation in the *Drosophila* visual system: A user’s guide to the dynamic morphology of the developing optic lobe. Dev Biol, 428, 1–24.

Nogi, T., Yasui, N., Mihara, E., Matsunaga, Y., Noda, M., Yamashita, N., Toyofuku, T., Uchiyama, S., Goshima, Y., Kumanogoh, A., et al (2010) Structural basis for Semaphorin signalling through the Plexin receptor. Nature, 467, 1123–1127.

Ozel, M. N., Simon, F., Jafari, S., Holguera, I., Chen, Y. C., Benhra, N., El-Danaf, R. N., Kapuralin, K., Malin, J. A., Konstantinides, N., et al (2021) Neuronal diversity and convergence in a visual system developmental atlas. Nature, 589, 88–U93.

Pascoe, H. G., Wang, Y. and Zhang, X. (2015) Structural mechanisms of Plexin signaling. Prog Biophys Mol Biol, 118, 161–168.

Pecot, Matthew Y., Tadros, W., Nern, A., Bader, M., Chen, Y. and Zipursky, S. L. (2013) Multiple interactions control synaptic layer specificity in the *Drosophila* visual system. Neuron, 77, 299–310.

Plazaola-Sasieta, H., Fernandez-Pineda, A., Zhu, Q. and Morey, M. (2017) Untangling the wiring of the *Drosophila* visual system: developmental principles and molecular strategies. J Neurogenet, 31, 231–249.

Rozbesky, D., Verhagen, M. G., Karia, D., Nagy, G. N., Alvarez, L., Robinson, R. A., Harlos, K., Padilla-Parra, S., Pasterkamp, R. J. and Jones, E. Y. (2020) Structural basis of Semaphorin-Plexin cis interaction. EMBO J, 39, e102926.

Sanes, J. R. and Zipursky, S. L. (2010) Design principles of insect and vertebrate visual systems. Neuron, 66, 15–36.

Schwabe, T., Borycz, J. A., Meinertzhagen, I. A. and Clandinin, T. R. (2014) Differential adhesion determines the organization of synaptic fascicles in the *Drosophila* visual system. Curr Biol, 24, 1304–1313.

Stedden, C. G., Menegas, W., Zajac, A. L., Williams, A. M., Cheng, S., Özkan, E. and Horne-Badovinac, S. (2019) Planar-polarized Semaphorin-5c and Plexin A promote the collective migration of epithelial cells in *Drosophila*. Current Biology, 29, 908–920.e906.

Sun, L. O., Brady, C. M., Cahill, H., Al-Khindi, T., Sakuta, H., Dhande, O. S., Noda, M., Huberman, A. D., Nathans, J. and Kolodkin, A. L. (2015) Functional assembly of accessory optic system circuitry critical for compensatory eye movements. Neuron, 86, 971–984.

Sun, L. O., Jiang, Z., Rivlin-Etzion, M., Hand, R., Brady, C. M., Matsuoka, R. L., Yau, K. W., Feller, M. B. and Kolodkin, A. L. (2013) On and off retinal circuit assembly by divergent molecular mechanisms. Science, 342, 1241974.

Suzuki, T., Hasegawa, E., Nakai, Y., Kaido, M., Takayama, R. and Sato, M. (2016) Formation of neuronal circuits by interactions between neuronal populations derived from different origins in the *Drosophila* visual center. Cell Reports, 15, 499–509.

Sweeney, L. B., Couto, A., Chou, Y. H., Berdnik, D., Dickson, B. J., Luo, L. and Komiyama, T. (2007) Temporal target restriction of olfactory receptor neurons by Semaphorin-1a/PlexinA-mediated axon-axon interactions. Neuron, 53, 185–200.

Tamagnone, L., Artigiani, S., Chen, H., He, Z., Ming, G.-l., Song, H.-j., Chedotal, A., Winberg, M. L., Goodman, C. S., Poo, M.-m., et al (1999) Plexins are a large family of receptors for transmembrane, secreted, and GPI-anchored Semaphorins in vertebrates. Cell, 99, 71–80.

Tayler, T. D., Robichaux, M. B. and Garrity, P. A. (2004) Compartmentalization of visual centers in the *Drosophila* brain requires Slit and Robo proteins. Development, 131, 5935–5945.

Terman, J. R., Mao, T. Y., Pasterkamp, R. J., Yu, H. H. and Kolodkin, A. L. (2002) MICALs, a family of conserved flavoprotein oxidoreductases, function in Plexin-mediated axonal repulsion. Cell, 109, 887–900.

Timofeev, K., Joly, W., Hadjieconomou, D. and Salecker, I. (2012) Localized Netrins act as positional cues to control layer-specific targeting of photoreceptor axons in *Drosophila*. Neuron, 75, 80–93.

Ting, C. Y., McQueen, P. G., Pandya, N., Lin, T. Y., Yang, M., Reddy, O. V., O’Connor, M. B., McAuliffe, M. and Lee, C. H. (2014) Photoreceptor-derived Activin promotes dendritic termination and restricts the receptive fields of first-order interneurons in *Drosophila*. Neuron, 81, 830–846.

Toyofuku, T., Zhang, H., Kumanogoh, A., Takegahara, N., Yabuki, M., Harada, K., Hori, M. and Kikutani, H. (2004) Guidance of myocardial patterning in cardiac development by Sema6D reverse signalling. Nature Cell Biology, 6, 1204–1211.

Trush, O., Liu, C., Han, X., Nakai, Y., Takayama, R., Murakawa, H., Carrillo, J. A., Takechi, H., Hakeda-Suzuki, S., Suzuki, T., et al (2019) N-Cadherin orchestrates self-organization of neurons within a columnar unit in the *Drosophila* medulla. J Neurosci, 39, 5861–5880.

Wernet, M. F., Labhart, T., Baumann, F., Mazzoni, E. O., Pichaud, F. and Desplan, C. (2003) Homothorax switches function of *Drosophila* photoreceptors from color to polarized light sensors. Cell, 115, 267–279.

Winberg, M. L., Noordermeer, J. N., Tamagnone, L., Comoglio, P. M., Spriggs, M. K., Tessier-Lavigne, M. and Goodman, C. S. (1998) Plexin A Is a neuronal Semaphorin receptor that controls axon guidance. Cell, 95, 903–916.

Winberg, M. L., Tamagnone, L., Bai, J. W., Comoglio, P. M., Montell, D. and Goodman, C. S. (2001) The transmembrane protein Off-track associates with Plexins and functions downstream of Semaphorin signaling during axon guidance. Neuron, 32, 53–62.

Wittes, J. and Schüpbach, T. (2019) A gene expression screen in *Drosophila melanogaster* identifies novel JAK/STAT and EGFR targets during oogenesis. G3 9, 47–60.

Xie, X., Tabuchi, M., Brown, M. P., Mitchell, S. P., Wu, M. N. and Kolodkin, A. L. (2017) The laminar organization of the *Drosophila* ellipsoid body is Semaphorin-dependent and prevents the formation of ectopic synaptic connections. eLife, 6, e25328.

Yamagata, M. and Sanes, J. R. (2008) Dscam and Sidekick proteins direct lamina-specific synaptic connections in vertebrate retina. Nature, 451, 465–469.

---- (2012) Expanding the Ig superfamily code for laminar specificity in retina: expression and role of Contactins. J Neurosci, 32, 14402–14414.

Yamagata, M., Weiner, J. A. and Sanes, J. R. (2002) Sidekicks: synaptic adhesion molecules that promote lamina-specific connectivity in the retina. Cell, 110, 649–660.

Yoo, S. K., Pascoe, H. G., Pereira, T., Kondo, S., Jacinto, A., Zhang, X. and Hariharan, I. K. (2016) Plexins function in epithelial repair in both *Drosophila* and zebrafish. Nature Comm, 7, 12282.

Yu, H.-H., Araj, H. H., Ralls, S. A. and Kolodkin, A. L. (1998) The transmembrane Semaphorin Sema I is required in *Drosophila* for embryonic motor and CNS axon guidance. Neuron, 20, 207–220.

Yu, L., Zhou, Y., Cheng, S. and Rao, Y. (2010) Plexin A-Semaphorin-1a reverse signaling regulates photoreceptor axon guidance in *Drosophila*. J. Neurosci., 30, 12151–12156.

