## Supplementary Figures and Legends for "Two distinct mechanisms of Plexin A function in *Drosophila* optic lobe lamination and morphogenesis"

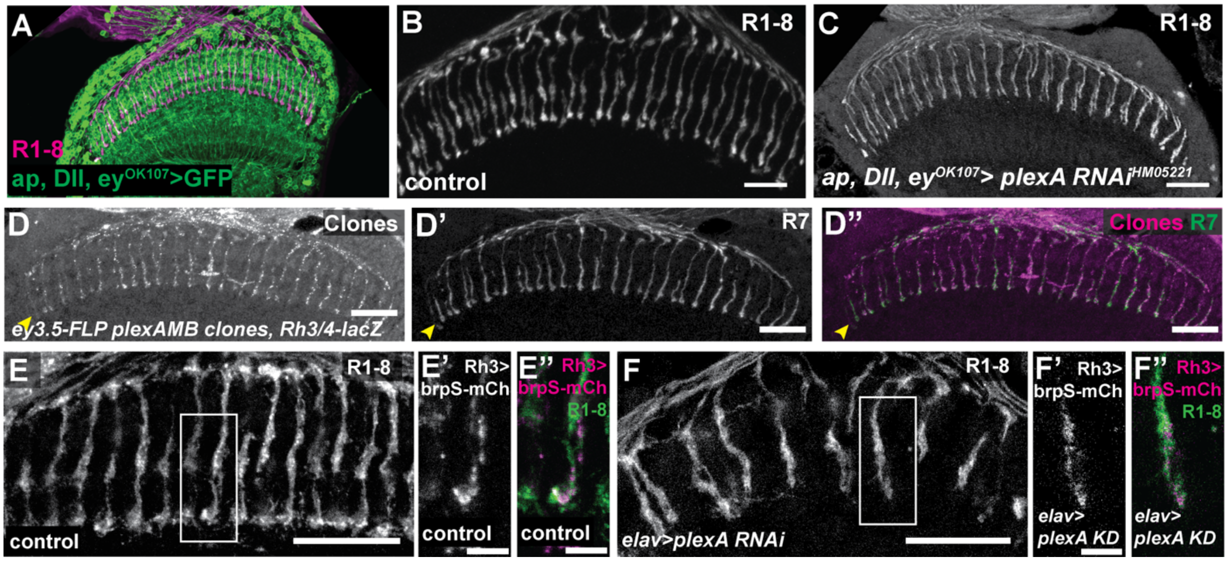


**Supplementary Figure 1. *plexA* is required for R7 photoreceptor targeting but not localization of synaptic components. (A)** *ap*-GAL4, *Dll*-GAL4, *ey*$^{OK107}$-GAL4 driving UAS-mCD8-GFP marks most medulla neurons (green). R1-8 are labeled with anti-Chp (magenta). **(B, C)** Adult head cryosections showing mistargeting of photoreceptors when *plexA* RNAi HM05221 is expressed in medulla neurons with *ap*-GAL4, *Dll*-GAL4, *ey*$^{OK107}$-GAL4 **(C)**, compared to control *w*$^{1118}$ **(B)**. Photoreceptors are marked with *gl-lacZ*. **(D)** Adult head cryosections showing large *plexA*$^{MB09499}$ homozygous clones generated in a *Minute* background using *ey*$^{3.5}$*-FLP*. R7 cells are marked with *Rh3/4-lacZ* (**D'**, green in **D''**), mutant photoreceptors are marked with *UAS-myr-Tomato* (**D**, magenta in **D''**). Arrowhead indicates the M6 layer of the medulla. **(E)** Adult head cryosections with R7 presynaptic sites labeled with *Rh3-LexA>LexAop-brpShort-mCherry* in **(E)** control and **(F)** *plexA* knockdown flies, where UAS-*plexA RNAi* HM05221 is expressed with the pan-neuronal driver *elav*-GAL4. R1-8 photoreceptors are labeled with anti-Chp (**E,** F; green in **E'** and **F'**); BrpS-mCherry is labeled with anti-dsRed (**E', F'**, magenta in **E''** and **F''**). Insets shown in **(E', E'', F', F'')** are enlargements of the boxes in **(E, F)**. **(A-F)** Scale bar = 20 μm; **(E'-F'')** scale bar = 5 μm.

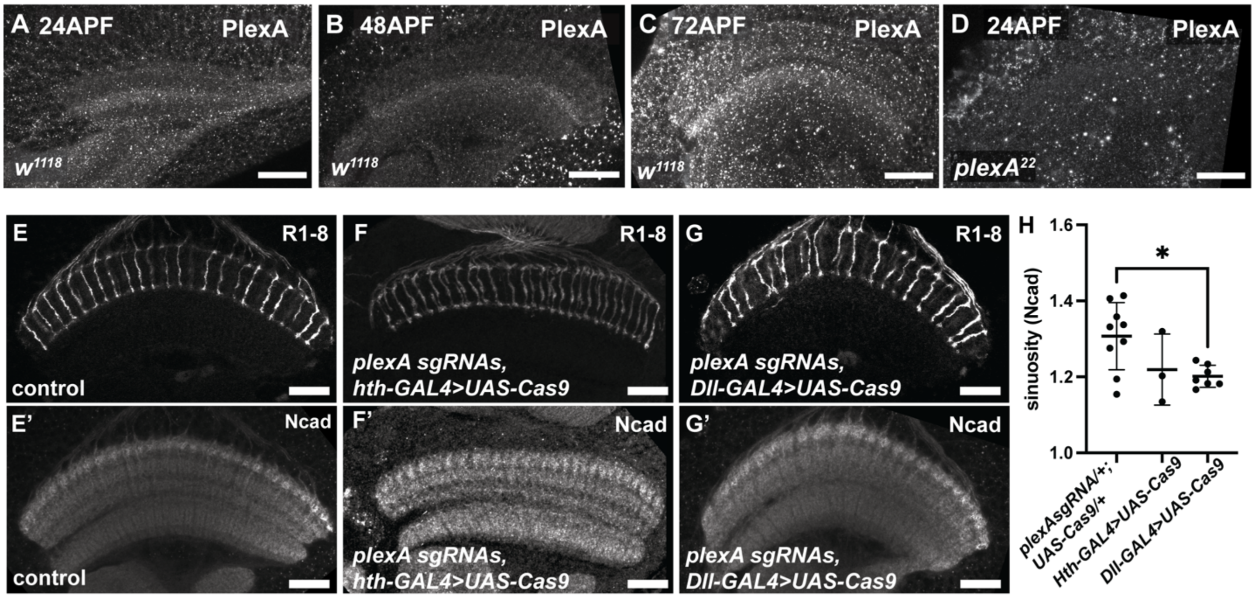


**Supplementary Figure 2. *plexA* is required in tangential neuron precursors for medulla layering. (A-D)** PlexA antibody staining of $w^{1118}$ 24 h APF **(A),** 48 h APF **(B)** and 72 h APF **(C)** and of $plexA^{22}$ 24 h APF **(D)** brains. **(E-G)** 72 h APF brains of control (*plexA sgRNA/+; UAS-Cas9/+*) **(E)** and somatic CRISPR *plexA* mutants generated by expressing *UAS-Cas9* with *hth-GAL4* **(F)** or *Dll-GAL4* **(G)** together with ubiquitously expressed *plexA* null sgRNAs in a background heterozygous for the $plexA^{MB}$ allele. R1-8 are labelled with anti-Chp **(E-G)** and the neuropil with Ncad **(E'- G')**. Scale bar = 20 µm. **(H)** Quantification of the sinuosity of Ncad intensity measurements taken along the height of the medulla. N = 9 control, 3 *hth-GAL4*, 7 *Dll-GAL4* samples. Means of each group were compared via one-way ANOVA with Tukey's multiple comparisons test. Error bars denote mean ± SD. *, $P \leq 0.05$.

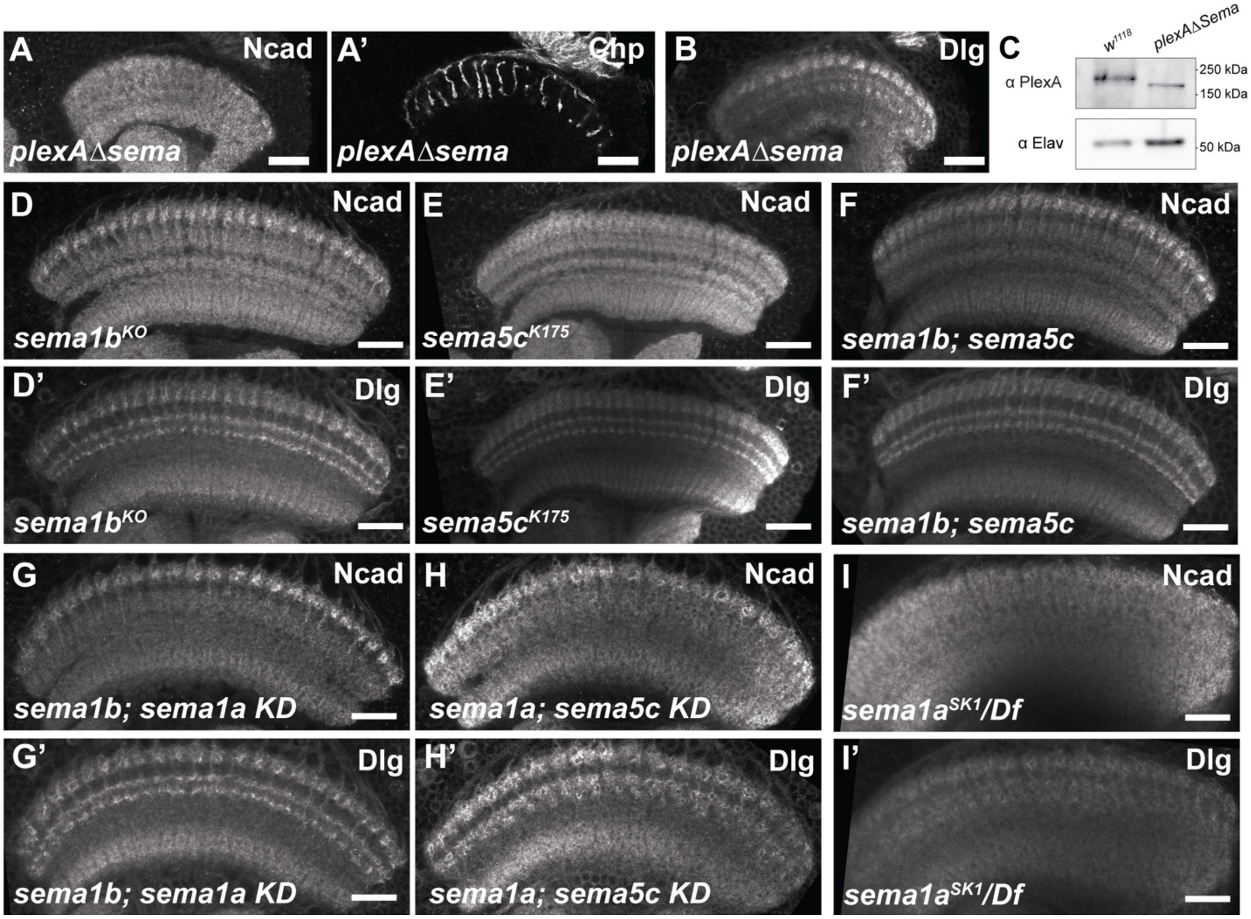


**Supplementary Figure 3. Sema1b and Sema5c do not have detectable effects on medulla layering. (A, B)** *plexAΔSema* mutant brains stained for Ncad **(A)**, Chp **(A')**, and Dlg **(B)**. **(C)** Western blot validation of the *plexAΔSema* allele demonstrating the presence of PlexAΔSema protein in third instar larval brains. **(D-I)** show 72 h APF pupal brains stained for Ncad **(D-I)** and Dlg **(D'-I')**. **(D)** *sema1b*$^{KO}$*;* **(E)**, *sema5c*$^{K175}$*;* **(F)**, *sema1b*$^{KO}$*; sema5c*$^{K175}$ double mutants; (**G)** *sema1a* RNAi expressed with nS*yb-GAL4* in a *sema1b*$^{KO}$ mutant background (compare to Fig. S4C); **(H)** *sema5c* RNAi expressed with nSyb-GAL4 in a *sema1a*$^{P1}$ mutant background; **(I)** *sema1a*$^{SK1}$*/Df(2L)Exel7039*. Scale bar = 20 μm.

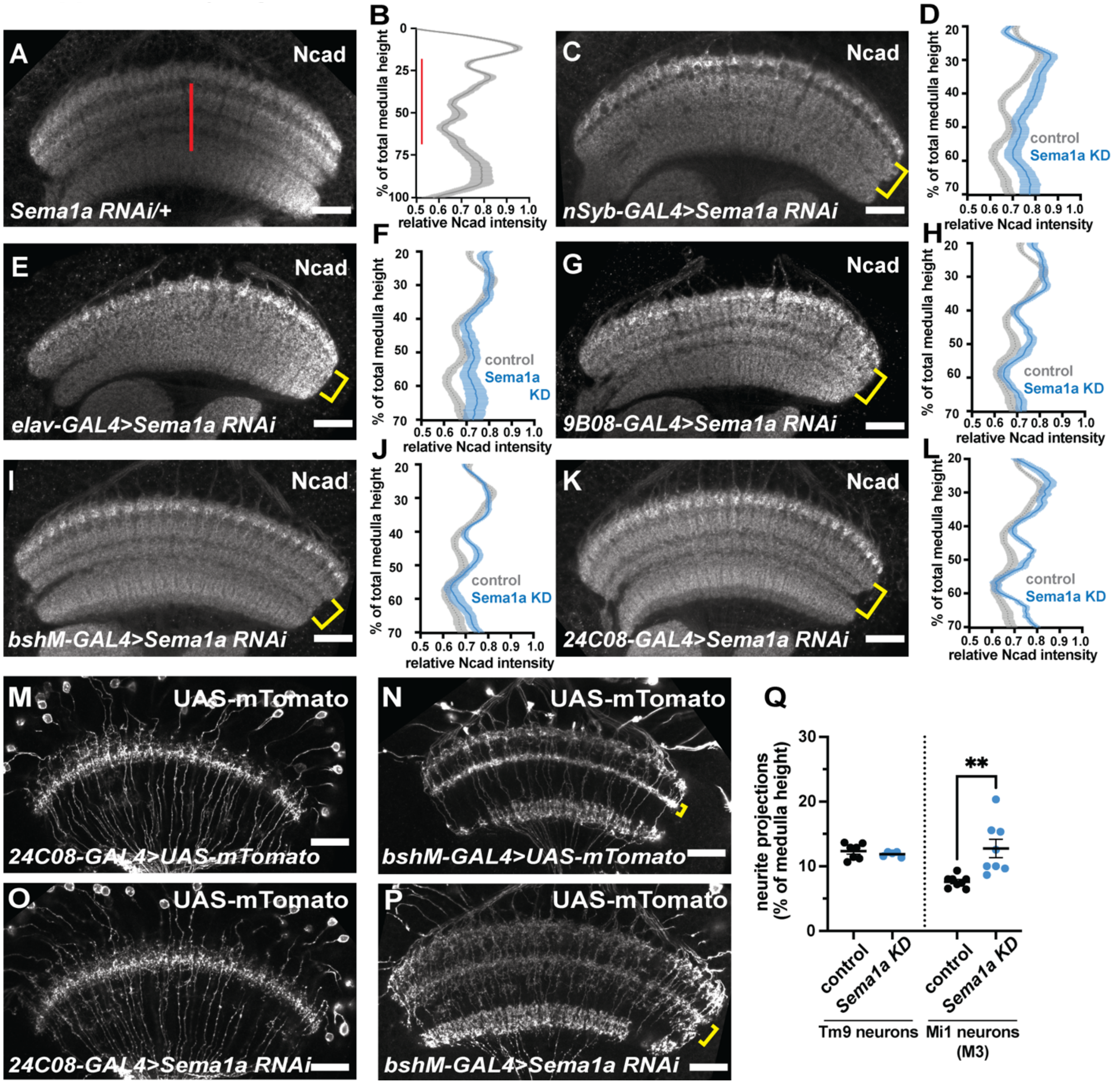


**Supplementary Figure 4. Lamina neurons are not the only source of Sema1a required for medulla layer formation. (A)** *UAS-Sema1a RNAi/+* control stained for Ncad and **(B)** quantification of Ncad intensity taken along the medulla height. Red line denotes the medial medulla (20-70% depth) quantified in **(C-L)**. **(C-L)** *sema1a* RNAi expressed with the pan-neuronal drivers *nSyb*-GAL4 **(C)** and *elav*-GAL4 **(E)**, the lamina neuron driver *GMR9B08-GAL4* **(G)**, the Mi1 neuron driver *bshM*-GAL4 **(I)**, and the Tm9 neuron driver *GMR24C08*-GAL4 **(K)**. Quantifications of Ncad intensity taken along the medial domain of the medulla are shown for *sema1a* RNAi expression with *nSyb*-GAL4 **(D)**, *elav*-GAL4 **(F),** *GMR9B08*-GAL4 **(H)**, *bshM*-GAL4 **(J)** and *GMR24C08*-GAL4 **(L)**. Yellow brackets denote medial medulla region of interest.

N = 15 control, 8 *nSyb-GAL4>Sema1a RNAi*, 6 *elav-GAL4>Sema1a RNAi*, 9 *9B08-GAL4>Sema1a RNAi*, 5 *24C08-GAL4>Sema1a RNAi*, 8 *bshM-GAL4>Sema1a RNAi* samples. **(M-P)** Visualization of control **(M, N)** and *sema1a* knockdown **(O, P)** Tm9 and Mi1 neurites using *UAS-myr-Tomato* driven by *24C08-GAL4* **(M, O)** or *bshM-GAL4* **(N, P). (Q)** Quantification of Tm9 and Mi1 (M3 layer, yellow brackets) neurite outgrowth as function of total medulla height. N = 7 control, 5 *sema1a KD* Tm9 samples; 8 Mi1 samples/genotype. Measurements were obtained in 72 h APF pupal brains in a horizontal plane and compared via unpaired t-test. Error bars denote mean ± SEM. **, P ≤ 0.01.

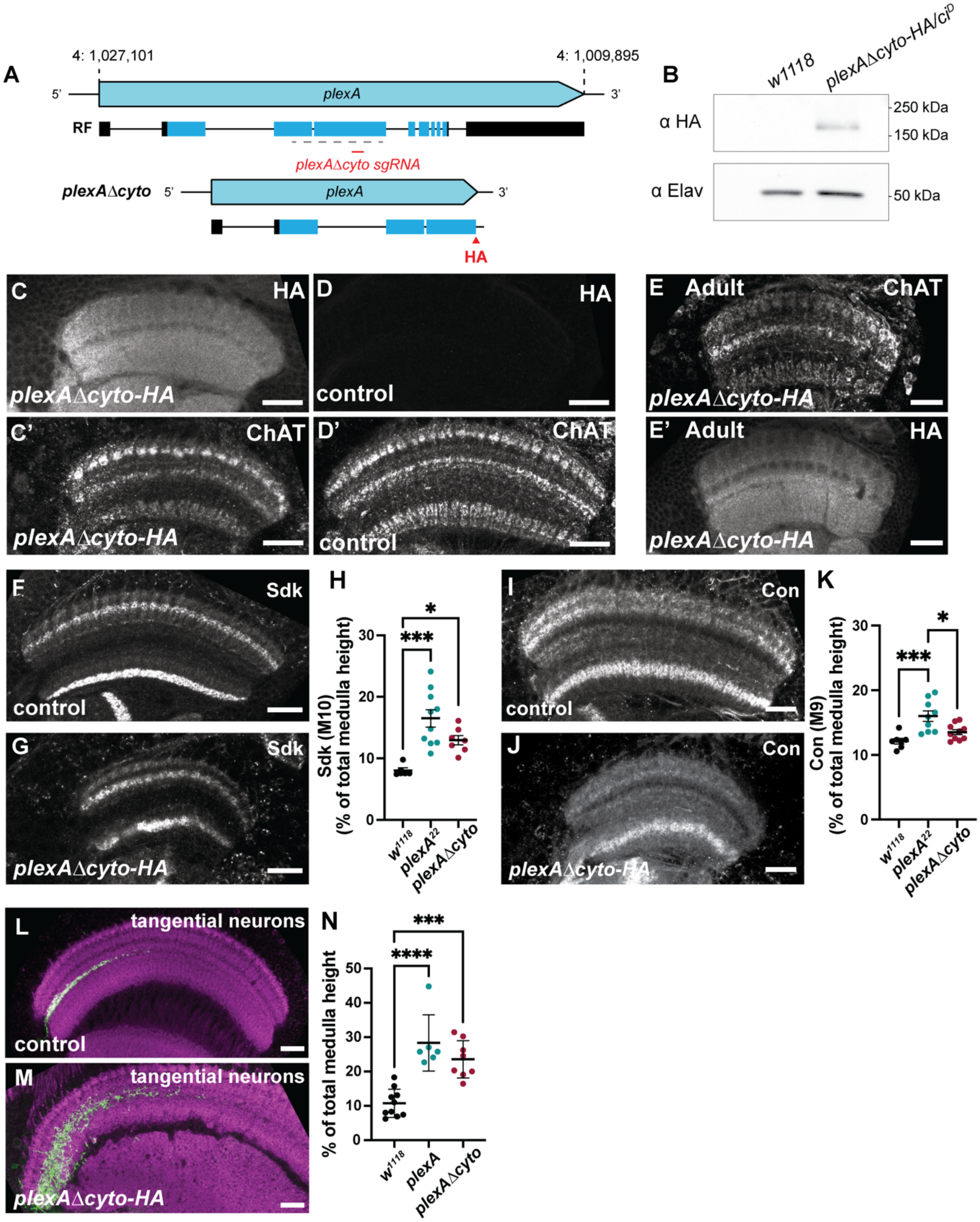


**Supplementary Figure 5. The cytoplasmic domain of PlexA is dispensable for medulla layering. (A)** Schematic depicting the generation of the *plexAΔcyto* mutant allele. Red line indicates location of *plexAΔcyto* sgRNA, grey dashed line indicates *plexA*$^{22}$ deletion. The resulting gene product and location of the HA tag are shown below. **(B)** Western blot validation of the *plexAΔcyto* allele demonstrating the presence of PlexAΔcyto-HA protein in third instar larval brains. **(C, D)** 72 h APF *plexAΔcyto-HA* mutant **(C)** and $w^{1118}$ control **(D)** 72 h APF brains stained with anti-HA **(C, D)** and anti-ChAT **(C', D')**. **(E)** Whole mount adult brain of *plexAΔcyto-HA* stained for ChAT **(E)** and HA **(E')**. **(F, G, I, J)** 72 h APF $w^{1118}$ control **(F, I)** and *plexAΔcyto-HA* mutant brains **(G, J)** stained with anti-Sdk **(F, G)** and anti-Con **(I, J). (H)** Quantification of width of Sidekick domain in M10 layer, plotted as percent of total medulla height. **(K)** Quantification of width of Connectin domain in M9 layer, plotted as percent of total medulla height. N = 5 WT, 10 $plexA^{22}$, 7 *plexAΔcyto* Sdk samples; 7 WT, 9 $plexA^{22}$, 10 *plexAΔcyto* Con samples. **(L, M)** Visualization of tangential neurons by *GMR35A02-GAL4* driving *UAS*-myrTomato in frontal optical sections of control **(L)** and *plexAΔcyto* mutant **(M)** brains. **(N)** Quantification of the extent of tangential neurite outgrowth. All quantifications are shown for $w^{1118}$ controls, $plexA^{22}/plexA^{MB}$ (*plexA*) and $plexA\Delta cyto/plexA^{22}$ (*plexAΔcyto*) mutants. N = 10 WT, 6 $plexA^{22}$, 8 *plexAΔcyto* samples. Means of each group were compared via one-way ANOVA with Tukey's multiple comparisons test. Error bars denote mean ± SEM. *, P ≤ 0.05; ***, P ≤ 0.001. Comparisons not shown were not significant.

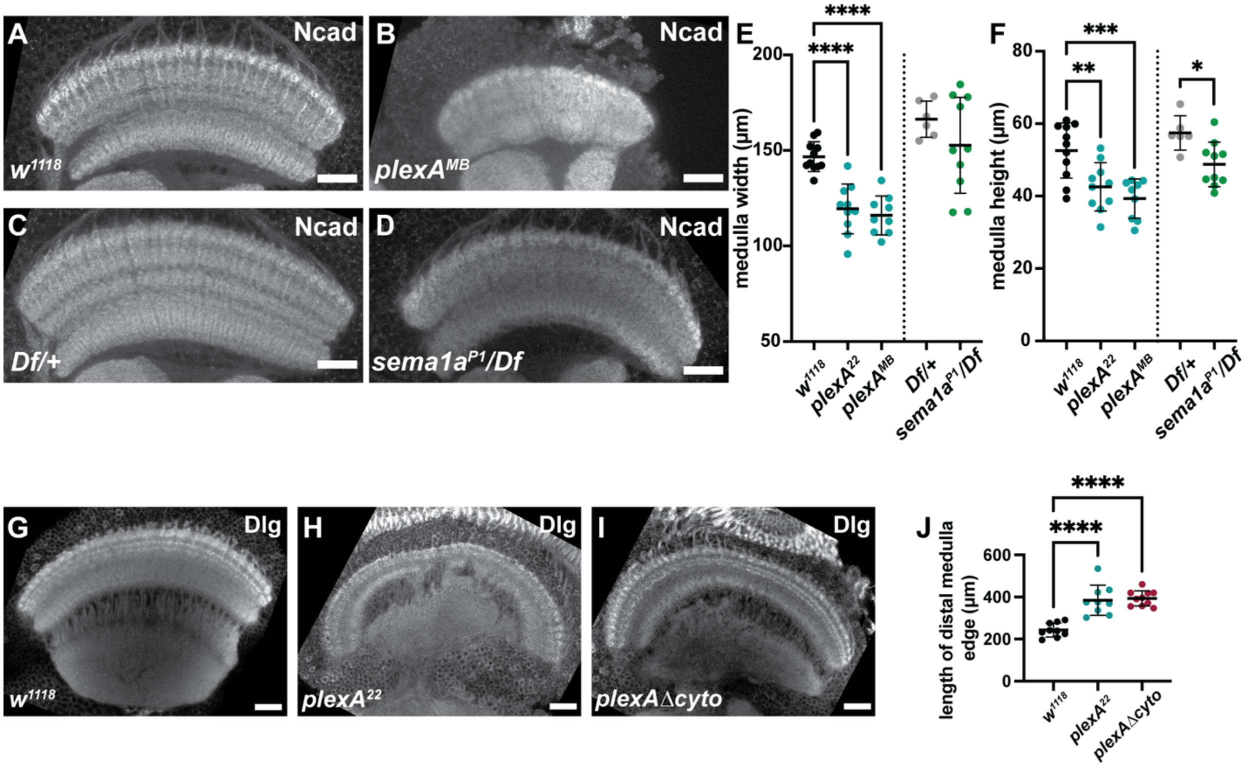


**Supplementary Figure 6. PlexA, but not Sema1a, is required to establish proper medulla morphology. (A-D)** Whole mount brains of *$w^{1118}$* **(A)**, *$plexA^{MB09499}$* **(B),** Df(2L)*Exel7039/+* **(C)** and *$sema1a^{P1}$/Df(2L)Exel7039* **(D)** 72 h APF pupae, stained with Ncad to visualize the neuropil. **(E, F)** Quantification of medulla width **(E)** and height **(F)** in these genotypes (*$plexA^{22}$* quantifications from Fig. 6J, K shown for comparison). Quantifications were done on 72 h APF pupal brains imaged in horizontal optical sections. N = 11 WT, 10 *$plexA^{22}$*, 9 *$plexA^{MB}$*, 6 *Df/+*, 10 *$sema1a^{P1}$/Df* samples. **(G-I)** Frontal optical sections of 72 h APF *$w^{1118}$* **(G)**, *$plexA^{22}$* **(H)**, and *plexAΔcyto* **(I)** pupal brains stained for Dlg. Ventral is to the left, dorsal is to the right. **(J)** Quantification of the distal perimeter of the medulla neuropil in frontal optical sections of these genotypes. Means of *plexA* and *plexAΔcyto* samples were compared with *$w^{1118}$* control samples via one-way ANOVA with Dunnetts multiple comparisons test; *Df/+* and *$sema1a^{P1}$/Df* were compared via unpaired t-test. N = 9 control, 9 *$plexA^{22}$*, 10 *plexAΔcyto* samples. Error bars denote mean ± SD. Scale bars = 20 µm. *, $P \leq 0.05$; **, $P \leq 0.01$; ***, $P \leq 0.001$; ****, $P \leq 0.0001$.
